# Calcium current modulation by the γ_1_ subunit depends on alternative splicing of Ca_V_1.1

**DOI:** 10.1101/2021.11.10.468074

**Authors:** Yousra El Ghaleb, Nadine J. Ortner, Wilfried Posch, Monica L. Fernández-Quintero, Wietske E. Tuinte, Stefania Monteleone, Henning J. Draheim, Klaus R. Liedl, Doris Wilflingseder, Jörg Striessnig, Petronel Tuluc, Bernhard E. Flucher, Marta Campiglio

## Abstract

The skeletal muscle voltage-gated calcium channel (Ca_V_1.1) primarily functions as voltage sensor for excitation-contraction coupling. Conversely, its ion-conducting function is modulated by multiple mechanisms within the pore-forming α_1S_ subunit and the auxiliary α_2_δ-1 and γ_1_ subunits. Particularly, developmentally regulated alternative splicing of exon 29, which inserts 19 amino acids in the extracellular IVS3-S4 loop of Ca_V_1.1a, greatly reduces the current density and shifts the voltage-dependence of activation to positive potentials outside the physiological range. We generated a new HEK293-cell line stably expressing α_2_δ-1, β_3_, and STAC3. When the adult (Ca_V_1.1a) and the embryonic (Ca_V_1.1e) splice variants were expressed in these cells, the difference in the voltage-dependence of activation observed in muscle cells was reproduced, but not the reduced current density of Ca_V_1.1a. Only when we further co-expressed the γ_1_ subunit, the current density of Ca_V_1.1a, but not of Ca_V_1.1e, was reduced by >50 %. In addition, γ_1_ caused a shift of the voltage-dependence of inactivation to negative voltages in both variants. Thus, the current-reducing effect of γ_1_, but not its effect on inactivation, is specifically dependent on the inclusion of exon 29 in Ca_V_1.1a. Molecular structure modeling revealed several direct ionic interactions between oppositely charged residues in the IVS3-S4 loop and the γ_1_ subunit. However, substitution of these residues by alanine, individually or in combination, did not abolish the γ_1_-dependent reduction of current density, suggesting that structural rearrangements of Ca_V_1.1a induced by inclusion of exon 29 allosterically empower the γ_1_ subunit to exert its inhibitory action on Ca_V_1.1 calcium currents.

**Summary:** El Ghaleb et al. analyzed the effects of the γ_1_ subunit on current properties and expression of the adult (Ca_V_1.1a) and embryonic (Ca_V_1.1e) calcium channel splice variants, demonstrating that γ_1_ reduces the current amplitude in a splicing-dependent manner.

## Introduction

Excitation-contraction (EC) coupling in skeletal muscle is initiated by action potentials that activate the voltage-gated calcium channel Ca_V_1.1 located in the transverse (T-) tubules. In adult skeletal muscle Ca_V_1.1 functions as voltage-sensor, which triggers the opening of the calcium release channel, the ryanodine receptor (RyR1), in the sarcoplasmic reticulum (SR) via protein-protein interactions, thus initiating muscle contraction (Rios and Brum 1987; Schneider and Chandler 1973). Because of the conformational coupling between Ca_V_1.1 and RyR1, Ca_V_1.1 currents are dispensable for skeletal muscle EC coupling (Armstrong, Bezanilla, and Horowicz 1972; Dayal et al. 2017). Accordingly, in mammals Ca_V_1.1 channels activate only upon strong, non-physiological membrane depolarization and conduct small and slowly activating currents (Tanabe et al. 1988). This is strikingly different in the embryonic splice variant (Ca_V_1.1e), which lacks 19 amino acids in the extracellular loop connecting segments S3 and S4 in the IV homologous repeat, due to alternative splicing excluding exon 29 (Tuluc et al. 2009). The embryonic Ca_V_1.1e isoform activates upon physiological membrane depolarization and conducts currents that are substantially larger in amplitude than those of the adult Ca_V_1.1a isoform.

Ca_V_1.1 is a multi-protein complex consisting of a pore-forming α_1_ subunit and several auxiliary proteins: the intracellular β_1a_, the glycosylphosphatidylinositol (GPI) anchored extracellular α_2_δ-1, and the transmembrane γ_1_ subunits (Curtis and Catterall 1984; Zamponi et al. 2015). While the β_1a_ subunit was shown to be essential for the functional expression of Ca_V_1.1 and for EC coupling (Gregg et al. 1996; Schredelseker et al. 2005), α_2_δ-1 and γ_1_ are dispensable for functional expression of Ca_V_1.1, but displayed an inhibitory effect on Ca_V_1.1 currents I(Freise et al. 2000; Obermair et al. 2005; Held et al. 2002; Ursu et al. 2001; Arikkath et al. 2003; Tuluc et al. 2009; Ahern et al. 2001). The α_2_δ-1 subunit slows down the kinetics of activation of Ca_V_1.1 currents, while the γ_1_ subunit reduces the current amplitude and shifts the voltage-dependence of inactivation.

All these studies were performed in skeletal muscle cells using a knockout or knockdown approach, since Ca_V_1.1 expresses poorly in mammalian non-muscle cells. Whereas co-expression of the auxiliary subunits β and α_2_δ is sufficient to support functional expression of all other high-voltage activated calcium channels (Singer et al. 1991; Lacerda et al. 1991; Zamponi et al. 2015), Ca_V_1.1 co-expression with these subunits does not yield functional currents in heterologous cell systems. Only recently, it was demonstrated that the skeletal muscle-specific adaptor protein STAC3 is essential for membrane expression and robust currents of Ca_V_1.1 in heterologous cells (Polster et al. 2015; Wu et al. 2018).

In the present study, we generated two HEK cell lines stably expressing the three subunits (STAC3, β_3_ and α_2_δ-1) necessary to support functional membrane expression of Ca_V_1.1. These cell lines provide a unique tool for analysis of wild-type and mutant Ca_V_1.1 channel currents and pharmacology in non-muscle cells. Interestingly, in contrast to what had been reported in myotubes, our current analysis of the adult and embryonic Ca_V_1.1 splice variants in the STAC3-HEK cell lines revealed no difference in current densities, while still displaying the typical differences in voltage dependence of activation. Because co-expression of γ_1_ inhibits the gating properties of Ca_V_1.1a calcium currents in skeletal muscle myotubes and in tsA201 cells (Polster et al. 2016; Freise et al. 2000; Ahern et al. 2001), and because the recent Ca_V_1.1 structure revealed an interaction of γ_1_ subunit with the IVS3-S4 loop of Ca_V_1.1a (Wu et al. 2016; Wu et al. 2015), we hypothesized that regulation of the gating properties of Ca_V_1.1 channels by the γ_1_ subunit occurs in a splice variant-dependent manner. Indeed, we found that co-expressed γ_1_ subunits selectively reduce the current density of the adult Ca_V_1.1a isoform, and not that of the embryonic Ca_V_1.1e isoform. In contrast, γ_1_ similarly shifted the voltage dependence of steady state inactivation to more negative voltages and increased Ca_V_1.1 membrane expression of both isoforms. Molecular modeling predicted several ionic interactions between the γ_1_ subunit and the IVS3-S4 linker of Ca_V_1.1a. However, site-directed mutagenesis of the putative ion-pair partners did not abolish γ_1_-dependent inhibition of the Ca_V_1.1a currents, suggesting an allosteric effect of exon 29 that is important for modulation of current density by the γ_1_ subunit.

## Material and methods

### Generation of stable cell lines

Two HEK293 cell lines stably expressing mouse STAC3 were generated using the Flp-In T-Rex system (Invitrogen). Host cells, already expressing human α_2_δ-1 and β_3_ subunits and containing a flippase recognition target (FRT) site, allowed the integration of STAC3 into the genome in a Flp recombinase-dependent manner. Briefly, the coding sequence of mouse STAC3 (Q8BZ71) was cloned into the pTO-HA-strepIII C GW FRT vector (containing a FRT site and a hygromycin resistance gene). To generate the cell line constitutively expressing STAC3 (**HEK-STAC3**), STAC3 expression was under the control of a CMV promoter. To generate the inducible STAC3 expression cell line (**HEK-TetOn-STAC3**), STAC3 expression was under the control of a CMV promoter with a tetracycline operator (TetOn) element. HEK293 host cells were transfected using the calcium phosphate method with either plasmid and a Flp recombinase-expressing vector (pOG44). Subsequently cells were selected with Hygromycin B (50 μg/ml; catalog #CP12.2, Lactan/Roth) and selection agents for the other subunits (see below), and single positive cell clones were cultured and characterized. The electrophysiological experiments for the characterization of the cell lines were carried out using the TetOn-STAC3 cell line (Fig. 3, 4, 6 and S1).

### Cell culture and transfection

Cells were cultured in DMEM (catalog #41966052, invitrogen) supplemented with 10% fetal bovine serum (F9665, Sigma), 2 mM l-glutamine (25030-032, Invitrogen), 10 U/ml penicillin-streptomycin (15140122, Invitrogen), and were maintained at 37°C in a humidified incubator with 5% CO^2^. For maintenance of the stable cell lines, selection agents for each subunit were applied regularly [STAC3, 50 μg/ml hygromycin B; β_3_, 500 μg/ml geneticin (10131035, Gibco); and α_2_δ-1, 15 μg/ml blasticidin S (A1113903, Gibco)].

For electrophysiological experiments, cells were plated on 35 mm dishes coated with poly-l-lysine (catalog #P2636, Sigma-Aldrich) and simultaneously transfected with 2 μg of DNA using Fugene HD (catalog #E2312, Promega), accordingly to the manufacturer instructions. For the Tet-On cell line, STAC3 expression was induced using 1 μg/ml doxycycline upon transfection (catalog #3072, Sigma-Aldrich), and cells were kept at 5% CO^2^ at 30°C. Cells were used for patch-clamp experiments 24–48 h after transfection/induction.

### Plasmids

Cloning procedures for GFP-Ca_V_1.1a and GFP-Ca_V_1.1e were previously described (Grabner, Dirksen, and Beam 1998; Tuluc et al. 2009).

Mouse γ_1_ was cloned from genomic cDNA from mouse soleus muscle. Primer sequences were selected according to Genebank NM-007582. Briefly, the cDNA of γ_1_ was amplified by PCR with a forward primer introducing a KpnI site upstream the starting codon (5’-ATATGGTACCATGTCACAGACCAAAACAGCGAAG-3’) and the reverse primer introducing a SalI site after the stop codon (5’-ATATGTCGACGCTAGTGCTCTGGCTCAGCGTCCATGCA-3’). The obtained PCR fragment, after KpnI/SalI digestion was inserted into the KpnI/XhoI digested pcDNA3 vector, yielding pcDNA3-γ_1_.

The 13-residue bungarotoxin binding site (BBS) was inserted in the IIS5-S6 loop of Ca_V_1.1a or Ca_V_1.1e at residue 593 by overlap extension PCR. Briefly the cDNA sequence of Ca_V_1.1 was amplified with overlapping primers in separate PCR reactions using GFP-Ca_V_1.1a as template. Primers used for the first fragment were: fw (5’-TACATGAGCTGGATCACG-3’) and rev (5’-GTAGGGCTCCAGGGAGCTCTCGTAGTATCTCCAGTGTCGCACTTCCGTGTCCTCGAAGTC -3’). Primers used for the second fragment were: fw (5’-TACGAGAGCTCCCTGGAGCCCTACCCTGACGTCACGTTCGAGGACACGGAAGTGCGACGC -3’) and rev (5’-GAACACGCACTGGACCACG -3’). The two separate PCR products were then used as template for a final PCR reaction with flanking primers to connect the nucleotide sequences. The resulting PCR fragment was then EcoRI/XhoI digested and inserted into the EcoRI/XhoI digested GFP-Ca_V_1.1a or GFP-Ca_V_1.1e, yielding GFP-Ca_V_1.1a-BBS or GFP-Ca_V_1.1e-BBS.

The R160A mutation was introduced by overlap extension PCR. Briefly the cDNA sequence of γ_1_ was amplified with overlapping primers mutating R160 into an alanine in separate PCR reactions using pcDNA3-γ_1_ as template. Primers used for the first fragment were: fw (5’-ATATGGTACCATGTCACAGACCAAAACAGCGAAG-3’) and rev (5’-CACCGACTGCGCCATGACCTCCACGGAGACGATGAG -3’). Primers used for the second fragment were: fw (5’-GAGGTCATGGCGCAGTCGGTGAAGCGTATGATTGAC-3’) and rev (5’-ATATGTCGACGCTAGTGCTCTGGCTCAGCGTCCATGCA-3’). The two separate PCR products were then used as template for a final PCR reaction with flanking primers to connect the nucleotide sequences. The resulting PCR fragment was then KpnI/SalI digested and inserted into the KpnI/XhoI digested pcDNA3 vector, yielding pcDNA3-γ_1_-R160A.

The K102A and E103A mutations were introduced by PCR. Briefly, the cDNA sequence of γ_1_ (nt 288-672) was amplified by PCR with a forward primer introducing the K102A and the E103A mutations downstream the EcoRI site and the reverse primer introducing an ApaI site after the stop codon. Primers used were: fw (5’-TGAATTCACCACTCAAGCGGCGTACAGCATCTCAGCAGCGGCCATT-3’) and rev (5’-AGAATAGGGCCCCCCCTCGACGCT-3’). The obtained PCR fragment, after EcoRI/ApaI digestion was inserted into the EcoRI/ApaI digested pcDNA3-γ_1_ vector, yielding pcDNA3-γ_1_-K102A-E103A. To combine the three mutations we introduced the K102A and the E103A mutations as described above, but using γ_1_-R160A as template for the PCR, yielding γ_1_-R160A-K102A-E103A (γ_1_-RK EAAA).

Sequence integrity of all newly generated constructs was confirmed by sequencing (MWG Biotech).

### RT-PCR

RNA was isolated from the three HEK293 cell lines after 48 h in culture using the RNeasy Protect Mini Kit (catalog #74124, Qiagen). After reverse transcription (Super-Script II reverse transcriptase, catalog #18064022, Invitrogen), the absolute number of transcripts in each sample was assessed by quantitative TaqMan PCR (Mm01159196_m1, Thermo Fisher Scientific), using a standard curve generated from known concentrations of a PCR product containing the target of the assay as described previously (Rufenach et al., 2020).

### Western Blotting

Proteins isolated from the three HEK cell lines were prepared as previously described (Campiglio and Flucher, 2017). Briefly, cells plated in 100 mm dishes were trypsinized after 48 h in culture. Cells were lysed in RIPA buffer with a pestle and left on ice for 30 minutes. The lysates were then centrifuged for 10 minutes. The protein concentration was determined using a BCA assay (catalog #23250, Pierce). 20 µg of protein samples were loaded on a NuPage gel (4-12% polyacrylamide, catalog #NP0321, Invitrogen) and separated by SDS-PAGE at 160 V. The protein samples were then transferred to a PVDF membrane at 25 V and 100 mA for 3 h at 4°C with a semidry-blotting system (Roth). The membrane was then cut and incubated with rabbit anti-STAC3 (1:2,000; catalog #20392-1, Proteintech, RRID:AB_10693618) or mouse anti-GAPDH (1:100,000; catalog #sc-32233, Santa Cruz Biotechnology, RRID:AB_627679) antibodies overnight at 4°C and then with HRP-conjugated secondary antibody (1:5,000, Pierce) for 1 h at room temperature. The chemiluminescent signal was developed with ECL Supersignal WestPico kit (catalog #34579, Thermo Scientific) and detected with ImageQuant LAS 4000.

### Immunocytochemistry

The three HEK cell lines were plated on poly-lysine coated coverslips and fixed in paraformaldehyde at RT after 2 days in culture. Fixed cells were incubated in 5% normal goat serum in PBS/BSA/Triton for 30 min. The rabbit anti-STAC3 antibody (1:2,000) was applied overnight at 4°C and detected with Alexa-594-conjugated secondary antibody. During the last washing step, cells were incubated with Hoechst dye to stain nuclei. Preparations were analyzed on an Axioimager microscope (Carl Zeiss, Inc) using a ×63, 1.4 NA objective. Images were recorded with a cooled CCD camera (SPOT; Diagnostic Instruments) and Metamorph image processing software (Universal Imaging, Corp.). Images were arranged in Adobe Photoshop 9 (Adobe Systems Inc.), and linear adjustments were performed to correct black level and contrast. To quantify the fluorescence intensity of the STAC3 staining, 14-bit gray scale images of the red (STAC3) and blue (Hoechst) channels were acquired for each cell line. A region of interest was manually traced around each cell in the STAC3 staining image, its intensity was recorded and background corrected using Metamorph. For each condition, between 15 and 31 cells were analyzed from each of three independent experiments.

### Labelling of cell surface Ca_V_1.1 channels with QD_655_

For cell-surface labeling a 13 amino acid high affinity bungarotoxin (BTX) binding site was inserted into Cav1.1a and Cav1.1e as described (Yang et al. 2010) and expressed in HEK-293 cells. 48h hours after transfection, cells were resuspended from 35 mm dishes with ice-cold PBS^++^ containing calcium and magnesium (pH 7.4, 0.9 mM CaCl_2_, 0.49 mM MgCl_2_), washed and incubated with 5 µM biotinylated α-bungarotoxin (catalog #B1196, Invitrogen) in PBS^++^/3% BSA in the dark for 1 h on ice. Cells were washed twice with PBS^++^/3% BSA and incubated with 10 nM streptavidin-conjugated quantum dots (QD_655_, catalog #Q10121MP, Invitrogen) in the dark for 1h on ice. Finally, cells were washed twice with PBS^++^/3% BSA and either assayed in flow cytometry or plated on poly-L-lysine coated coverslips and imaged.

### Microscopy

Cells were imaged in Tyrode’s physiological solution using a 63x, 1.4 NA objective Axioimager microscope (Carl Zeiss). 14-bit images were recorded with a cooled CCD camera (SPOT, Diagnostic Instruments) and Metaview image processing software (Universal Imaging). Image composites were arranged in Adobe photoshop CS6.

### Multiparameter flow cytometry

Labeled cells were counted by flow cytometry using a BD FACSVerse analyzer (Becton Dickinson, Franklin Lakes, NJ). For flow cytometric analyses, labeled cells were counted and analyzed using BD FACSuite v1.0.6 and BD FACS Diva v9.0 software (Becton Dickinson, Franklin Lakes, NJ). Cells expressing GFP were excited at 488nm and red signal was excited at 633nm. For each set of experiments untransfected or unlabeled cells, as well as single color controls, were used to adjust threshold values and setting were then used for analyzing all samples.

### Electrophysiology

Calcium currents in HEK cells were recorded with the whole-cell patch-clamp technique in voltage-clamp mode using an Axopatch 200A amplifier (Axon Instruments). Patch pipettes (borosilicate glass; Science Products) had resistances between 1.8 and 4.0 MΩ when filled with (mM) 135 CsCl, 1 MgCl_2_, 10 HEPES, 10 EGTA and 4 ATP-Na2 (pH 7.4 with CsOH). The extracellular bath solution contained (mM) 15 CaCl_2_, 150 choline-chloride, 10 HEPES, and 1 Mg-Cl_2_ (pH 7.4 with CsOH). Data acquisition and command potentials were controlled by pCLAMP software (Clampex version 10.2; Axon Instruments); analysis was performed using Clampfit 10.7 (Axon Instruments) and SigmaPlot 12.0 (SPSS Science) software. The current-voltage dependence of activation was determined using 300 or 500 ms long square pulses to various test potentials (holding potential -80 mV) and curves were fitted according to:

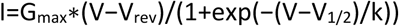

where G_max_ is the maximum conductance, V_rev_ is the extrapolated reversal potential, V_1/2_ is the potential for half maximal activation, and k is the slope. The conductance was calculated using G = (− I * 1000)/(V_rev_ − V), and its voltage dependence was fitted according to a Boltzmann distribution:

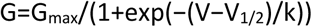

Steady-state inactivation was calculated as the ratio between two current amplitudes elicited by 200 ms pulses to V_max_ separated by a 15 second conditioning pulse to various test potentials (sweep start-to-start interval 30 seconds; see inset Fig. 4A). Steady-state inactivation curves were fitted using a modified Boltzmann equation:

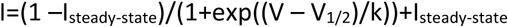

where V_1/2_ is the half-maximal inactivation voltage and k is the inactivation slope factor.

All experimental groups were analyzed in transiently transfected cells from at least three independent cell passages. The means, standard errors (SEM), and p-values were calculated using the Student’s t-test, 2-tailed, with significance criteria p < 0.05 *, p < 0.01 **, p < 0.001 *** and p < 0.0001 ****. P-values of the experiments where more than 2 groups are compared to each other were calculated using the ANOVA and Tukey’s or Sidak’s posthoc test.

### Structure modeling

The complex structures of both splice variants of the human α1-subunit (Ca_V_1.1e and Ca_V_1.1a) and the γ_1_-subunit were modelled based on the rabbit cryo-electron microscopy (EM) structure of Ca_V_1.1 in the inactivated state, with voltage sensors in the ‘up’ conformation and a closed intracellular gate (PDB accession code: 5GJV) (Wu et al. 2016). Homology modelling has been performed using MOE (Molecular Operating Environment, version 2018.08, Molecular Computing Group Inc., Montreal, Canada). Additionally, ab initio Rosetta modelling was used to generate structures for loops that were not resolved in the original Ca_V_1.1 α1-subunit and γ_1_-subunit template (Rohl et al. 2004). The structures for the putative mutants were derived from both WT splice variant models by replacing the mutated residue and carrying out a local energy minimization using MOE. The C-terminal and N-terminal parts of each domain were capped with acetylamide (ACE) and N-methylamide to avoid perturbations by free charged functional groups. The structure model was embedded in a plasma membrane consisting of POPC (1-palmitoyl-2-oleoyl-sn-glycero-3-phosphocholine) and cholesterol in a 3:1 ratio, using the CHARMM-GUI Membrane Builder (Lee et al. 2019; Jo et al. 2009). Water molecules and 0.15 M KCl were included in the simulation box. Energy minimizations of Ca_V_1.1e and Ca_V_1.1a WT and mutant structures in the membrane environment were performed. The topology was generated with the LEaP tool of the AmberTools18 (Case et al. 2008), using force fields for proteins and lipids, ff14SBonlysc and Lipid14 (Dickson et al. 2014), respectively. The structure models were heated from 0 to 300 K in two steps, keeping the lipids fixed, and then equilibrated over 1 ns. Then molecular dynamics simulations were performed for 10 ns, with time steps of 2 fs, at 300 K and in anisotropic pressure scaling conditions. Van der Waals and short-range electrostatic interactions were cut off at 10 Å, whereas long-range electrostatics were calculated by the Particle Mesh Ewald (PME) method (Salomon-Ferrer et al. 2013). As the extracellular loop 1 was not resolved in the Cryo-EM structure, we modelled 100 loop structures with Rosetta ab initio modelling (Rohl et al. 2004). By clustering on the loops using a RMSD distance criterion of 2 Å, we obtained 10 clusters. These 10 clusters were carefully evaluated and the energetically most favorable two cluster representatives, which formed interactions with the S3-S4 loop of VSD IV (exon 29), were considered for further minimizations in the membrane environment. MOE and Pymol was used to visualize the key interactions and point out differences in structure models (The PyMOL Molecular Graphics System, Version 2.0 Schrödinger, LLC.).

### Online supplementary material

Fig. S1 shows the activation and inactivation kinetics analysis pertaining to Fig. 3. Table S1 summarizes electrophysiological parameters pertaining to Fig.2. Table S2 summarizes electrophysiological parameters pertaining to Fig.3 and Fig. S1. Table S3 summarizes electrophysiological parameters pertaining to Fig.6.

## Results

### Generation of two HEK cell lines expressing β_3_, α_2_δ-1 and STAC3

In order to generate HEK293 cell lines that could reliably support Ca_V_1.1 expression, we inserted STAC3 into the genome of a host cell line already stably expressing α_2_δ-1 and β_3_ using the Flp-In T-Rex system. We generated two cell lines: one in which the expression of STAC3 was constitutive (HEK-STAC3) and one in which the expression of STAC3 was doxycycline (DOX) inducible (HEK-TetOn-STAC3). While the parental HEK cell line showed neither STAC3 mRNA nor protein expression, the selected clone of the constitutive HEK-STAC3 cell line strongly expressed STAC3 (Fig 1). As expected, without DOX induction the selected clone of the inducible HEK-TetOn-STAC3 cell line showed only weak basal STAC3 mRNA and protein expression. However, 24 h after the beginning of DOX induction, STAC3 expression levels were strongly increased and comparable to those of the constitutive HEK-STAC3 cell line (Fig. 1).

**Figure 1.**
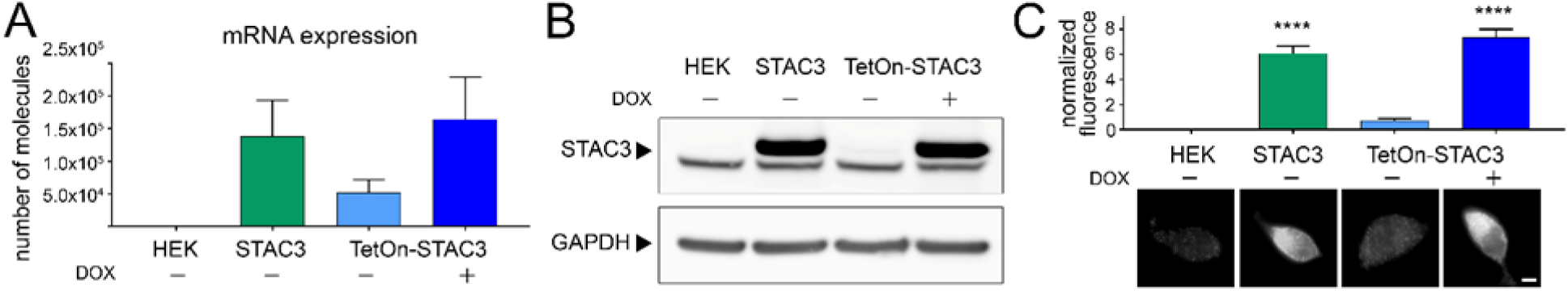
Both the constitutive and the inducible STAC3-HEK cell lines robustly express STAC3. **(A)** STAC3 mRNA transcription levels in the host (HEK), the constitutive (STAC3), and the inducible cell line (TetOn-STAC3), before and after doxycycline (DOX) treatment, assessed by TaqMan quantitative PCR. Mean values of three replicates. **(B)** Western blot analysis with anti-STAC3 antibody indicated that STAC3 is substantially expressed by the constitutive and the inducible cell lines (treated with DOX), while it is absent from the host cell line (HEK). Without DOX the inducible cell line shows very low basal expression. A non-specific band present in all samples migrates slightly fast than STAC3. One representative experiment of three is shown. **(C)** Quantification of STAC3 staining intensity in the host (HEK), the constitutive (STAC3), and the inducible cell line (TetOn-STAC3), before and after DOX treatment reveals a strong STAC3 expression in both STAC3 and TetOn-STAC3 cell lines. Scale bar, 2 µm. ANOVA, F(3,169)=67.72; P<0.0001; Tukey post hoc analysis ****P<0.0001.

We then analyzed the ability of the cell lines to support the expression of functional Ca_V_1.1 currents by transient transfection with the adult Ca_V_1.1a or the embryonic Ca_V_1.1e isoforms. The two Ca_V_1.1 isoforms differ in the length of the linker connecting helices S3 and S4 of the fourth homologous repeat, with the embryonic isoform skipping exon 29 and lacking 19 amino acids. Although both isoforms support skeletal muscle EC coupling, they display very different current properties when expressed in dysgenic (Ca_V_1.1-null) myotubes. In contrast to the adult Ca_V_1.1a isoform, the embryonic Ca_V_1.1e splice variant activates at more hyperpolarizing potentials and conducts calcium currents that are several-fold larger than those of Ca_V_1.1a (Tuluc et al. 2009). Our experiments show that both the constitutive (HEK-STAC3) and the inducible (HEK-TetOn-STAC3) cell lines efficiently supported functional expression of both the adult and the embryonic Ca_V_1.1 variants (Fig. 2A-B and 2E-F, Table S1). More interestingly, while the two Ca_V_1.1 splice variants displayed the expected difference in the V_½_ of activation (Fig. 2C-D and 2G-H, Table S1), the expected smaller current density in Ca_V_1.1a was not observed in the two STAC3-HEK cell lines (Fig. 2A-B, and 2E-F, Table S1).

**Figure 2.**
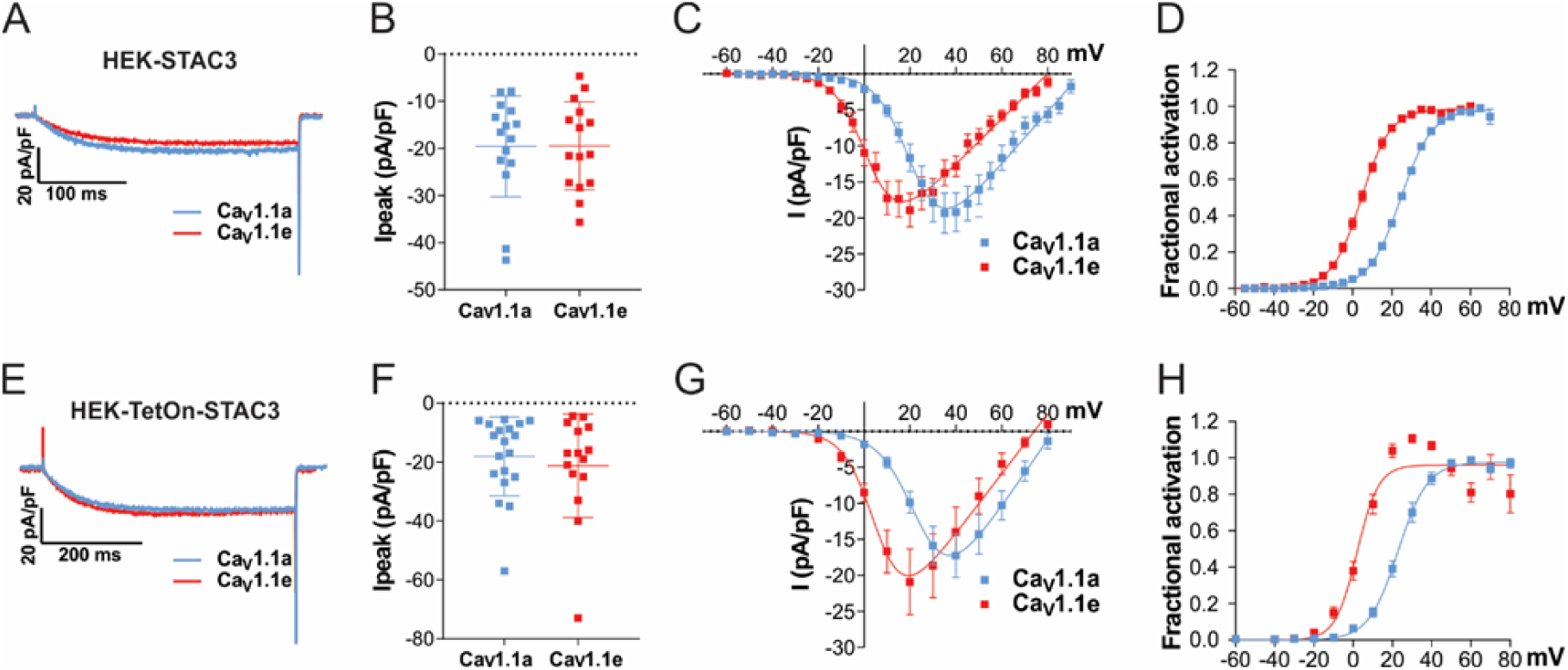
Exclusion of exon 29 in Ca_V_1.1e shifts the voltage dependence of activation to more negative voltages but does not affect current density in either of the two STAC3-HEK cell lines. **(A-D)** Current properties of Ca_V_1.1a (blue, n=15) compared to Ca_V_1.1e (red, n=15) in the HEK-STAC3 cell line. **(E-H)** Current properties of Ca_V_1.1a (blue, n=19) compared to Ca_V_1.1e (red, n=15) in the inducible cell line HEK-TetOn-STAC3 treated with DOX. **(A, E)** Exemplary current traces at V_max_ show similar activation kinetics of the Ca_V_1.1a and Ca_V_1.1e variants and no difference in the peak current density (I_peak_; peak current normalized to the cell size) in both the HEK-STAC3 **(B)** and HEK-TetOn-STAC3 **(F)** cell lines (p=0.94 and p=0.56, respectively). **(C, G)** The current-voltage relationship and **(D, H)** the normalized steady-stateactivation curves show that exclusion of exon 29 (in Ca_V_1.1e) results in a 20.6 mV and 21.1 mV left shift of activation when expressed in the HEK-STAC3 and HEK-TetOn-STAC3 cell line, respectively. Mean±SEM; P-values calculated with Student’s t-test (see Table S1 for parameters and statistics).

We reasoned that some factor is missing in HEK cells that specifically mediates the splicing-dependent effect on the current amplitude in muscle cells. Because in muscle the specific function of exon 29 is to curtail the calcium currents and in our STAC3-HEK cells the currents were equally large, the missing factor might be a muscle-specific protein capable of diminishing Ca_V_1.1 currents specifically in the adult splice variant. The only Ca_V_1.1 subunit not present in our expression system is the γ_1_ subunit. Moreover, the γ_1_ subunit acts as a negative regulator of Ca_V_1.1 currents both in skeletal muscle and in tsA201 cells (Freise et al. 2000; Ahern et al. 2001; Andronache et al. 2007; Polster et al. 2016) and its expression is restricted to skeletal muscle (Biel et al. 1991; Jay et al. 1990). Therefore, we inferred that the γ_1_ subunit may be the missing factor selectively reducing the currents of Ca_V_1.1a and not those of Ca_V_1.1e. This notion was further supported by the fact that cryo-EM structures of Ca_V_1.1 predicted an interaction of the γ_1_ subunit with the Ca_V_1.1 IVS3-S4 region, exactly the site containing the alternatively spliced exon 29 (Wu et al. 2016; Wu et al. 2015).

### The γ_1_ subunit selectively reduces the current density of Ca_V_1.1a but not that of Ca_V_1.1e

To test this hypothesis, we measured the calcium current density of Ca_V_1.1a and Ca_V_1.1e in the presence and the absence of γ_1_ in one of the newly established cell lines (HEK-TetOn-STAC3). As previously reported (Polster et al. 2016; Freise et al. 2000), the presence of γ_1_ significantly reduced Ca_V_1.1a current amplitudes, with no significant effect on the voltage dependence of activation (Fig. 3A-D and Table S2). The activation kinetics were unaltered by co-expression of the γ_1_ subunit (Fig. S1A-D, Table S2), in agreement to what had been observed in myotubes (Freise et al. 2000), but contrary to what was previously reported in tsA201 cells (Polster et al. 2016). More importantly, as hypothesized, in contrast to Ca_V_1.1a, co-expression of γ_1_ had no effect on the current density of Ca_V_1.1e (Fig. 3E-H, Table S2), suggesting that the inclusion of the 19 amino acids encoded in exon 29 is essential for suppression of the Ca_V_1.1 current by the γ_1_ subunit.

**Figure 3.**
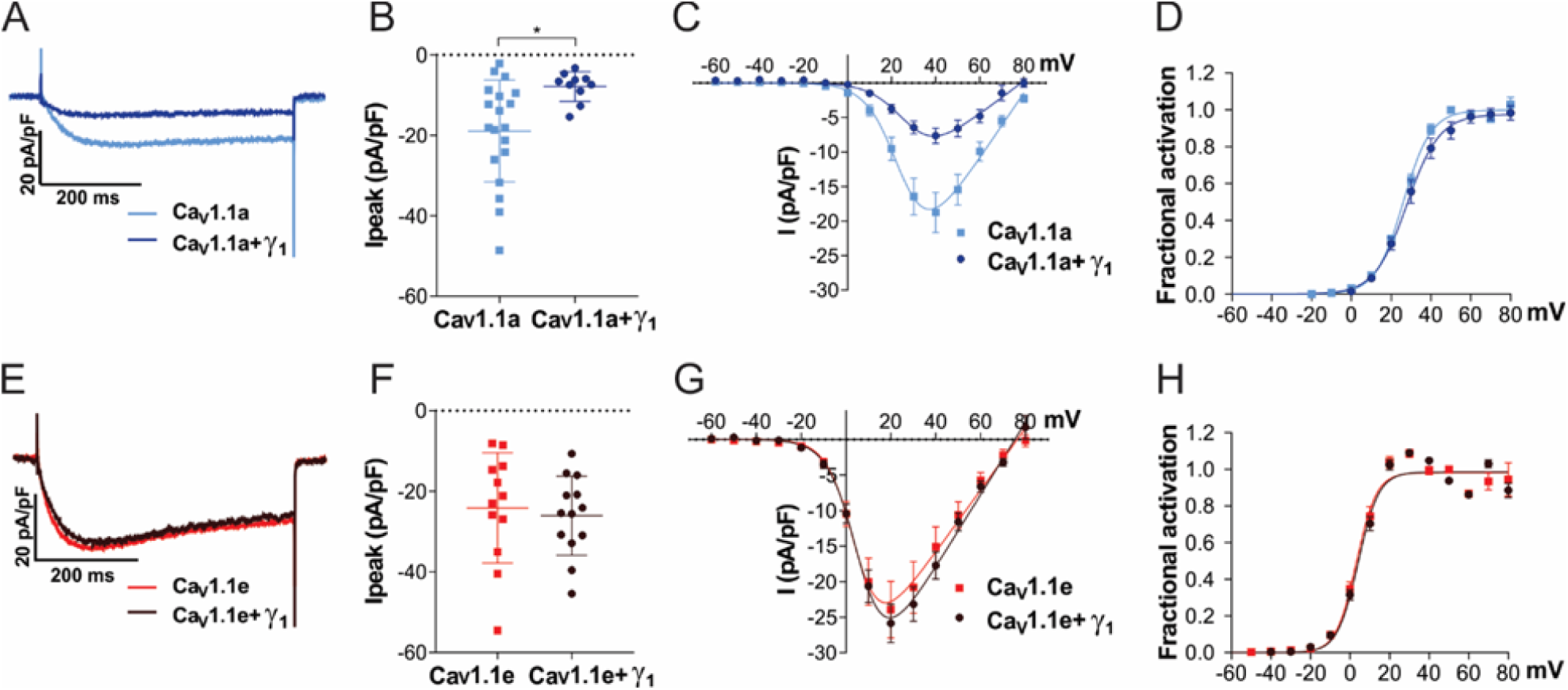
Co-expression of γ_1_ reduces the current density in Ca_V_1.1a but not in Ca_V_1.1e. **(A-D)** Current properties of the adult splice variant Ca_V_1.1a (blue, n=19) compared to Ca_V_1.1a co-expressed with γ_1_ (Ca_V_1.1a + γ_1_, dark blue, n=10). **(E-H)** Current properties of the embryonic splice variant Ca_V_1.1e (red, n=12) compared to Ca_V_1.1e + γ_1_ (dark red, n=13). **(A)** Exemplary current traces at V_max_ and **(B)** the scatter plot of the peak current density (I_peak_) show a significant reduction (p=0.012) when co-expressing γ_1_ with Ca_V_1.1a. On the contrary, when co-expressing γ_1_ with Ca_V_1.1e **(E-F)**, no difference in current density was observed (p=0.69). **(C, G)** The current-voltage relationship and **(D, H)** the fractional steady-state activation curves show no effect of γ_1_ on the voltage dependence of activation when co-expressed with Ca_V_1.1a or Ca_V_1.1e. Mean±SEM; *P*-values calculated with Student’s t-test. * *P<*0.05 (for parameters and statistics see Table S2).

### The γ_1_ subunit shifts the steady state inactivation to more negative potentials and decreases the window current of both Ca_V_1.1 isoforms

The γ_1_ subunit inhibits Ca_V_1.1 currents not only by decreasing the current amplitude, but also by limiting Ca_V_1.1 window current, i.e. the overlapping area between activation and inactivation curves. In fact, previous studies demonstrated that, in the presence of γ_1_, the voltage-dependence of inactivation shifted toward more negative potentials, while the voltage-dependence of activation remained unaltered (Ahern et al. 2001; Freise et al. 2000; Held et al. 2002; Ursu et al. 2004).

To determine if this γ_1_ effect on Ca_V_1.1 currents is also restricted to the adult Ca_V_1.1a isoform, we performed a steady state inactivation protocol comparing the current size of test pulses before and after 15 s conditioning pre-pulses at incrementally increasing potentials (inset in Fig. 4A). The normalized steady-state inactivation is plotted as a function of the pre-pulse potential. As previously demonstrated, co-expression of the γ_1_ subunit resulted in a robust left-shift in the voltage-dependence of inactivation of the adult Ca_V_1.1a isoform (Fig. 4A). In the presence of γ_1_, the half-maximal inactivation potential was shifted by 18.5 mV toward more hyperpolarizing potentials (Fig. 4B, Table S2), and inactivation was more complete compared to Ca_V_1.1a alone (Table S2). Consequently, the window current was decreased (even without considering the reduced current density in this voltage range) and peaked at ≈ 20 mV, as compared to ≈ 30 mV Ca_V_1.1a without γ_1_ (Fig 4C-D, Table S2).

**Figure 4.**
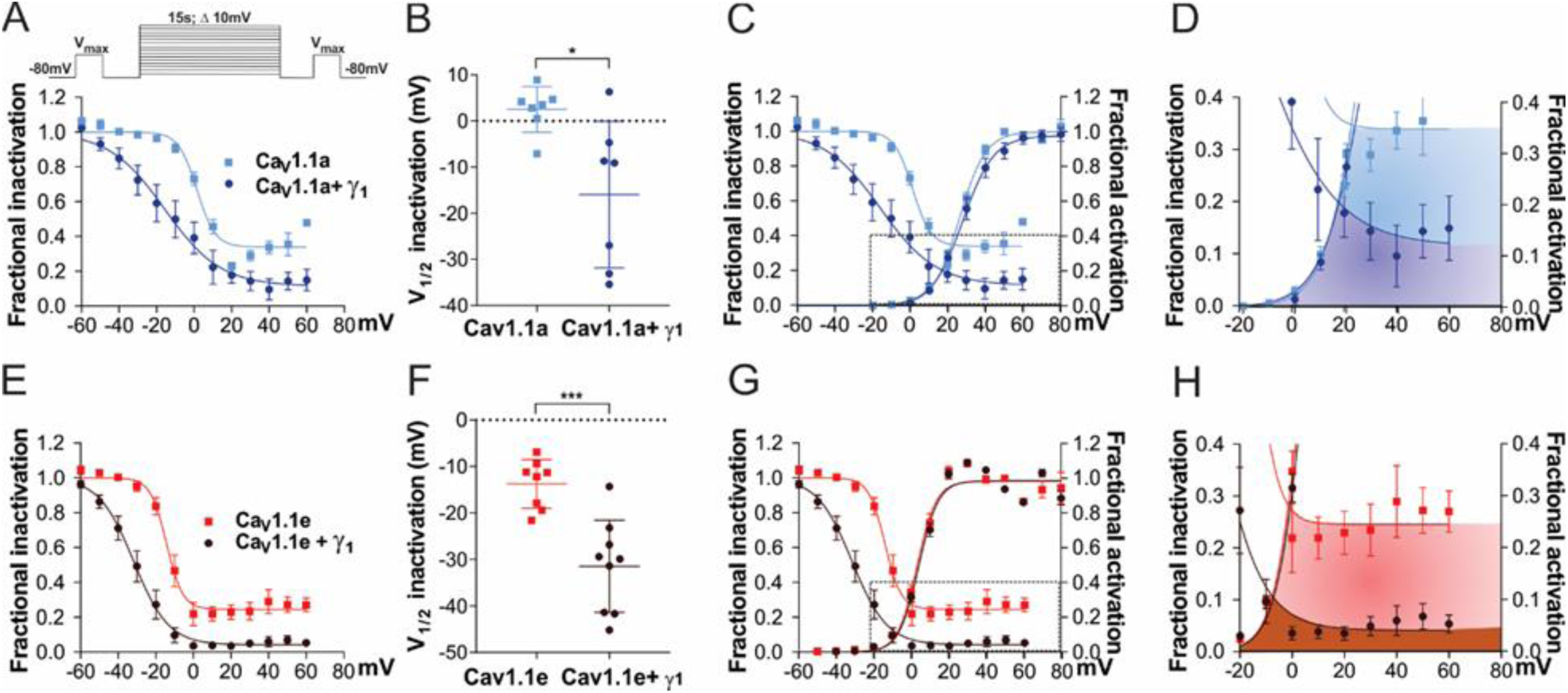
γ_1_ left shifts the steady state inactivation and reduces the window currents in both Ca_V_1.1a and Ca_V_1.1e. **(A-D)** Steady-state inactivation and window currents of Ca_V_1.1a (blue, n=7) compared to Ca_V_1.1a + γ_1_ (dark blue, n=7); (**E-H)** the same for Ca_V_1.1e (red, n=8) and Ca_V_1.1e + γ_1_ (dark red, n=9). **(A** and **B, E** and **F)** Fractional inactivation curves and scatter plot of V_½_ of inactivation show that, compared to Ca_V_1.1a and Ca_V_1.1e without γ_1_ expression, the voltage-dependence of inactivation is left-shifted in Ca_V_1.1a + γ_1_ (18.5 mV, p=0.013) and Ca_V_1.1e + γ_1_ (17.8 mV, p<0.001). The inset in **(A)** shows the steady state inactivation protocol. **(C, G)** Superposition of the fractional activation and inactivation curves (same as in panels A and E) shows that, while the activation of Ca_V_1.1a + γ_1_ and Ca_V_1.1e + γ_1_ is not shifted, the left shift in inactivation results in greatly decreased window currents. **(D, H)** The enlarged areas indicated by the frames (in **C** and **G)** show the size and voltage-range of the window currents for Ca_V_1.1a (shaded in blue), Ca_V_1.1a + γ_1_ (dark blue), Ca_V_1.1e (red) and Ca_V_1.1e + γ_1_ (dark red). Mean±SEM; *P*-values calculated with Student’s t-test. * *P<*0.05, *** *P*<0.001.

Surprisingly, these γ_1_ effects were recapitulated with the embryonic Ca_V_1.1e isoform. In the presence of the γ_1_ subunit, the half maximal inactivation potential was shifted to hyperpolarizing potentials by 17.8 mV and steady-state inactivation was almost complete (Fig-4E-F, Table S2). Because this left-shift of inactivation was not accompanied by a similar shift of V_½_ of activation the window current was decreased by several-fold and peaked at ≈ -10 mV, as compared to ≈ 0 mV in Ca_V_1.1e without γ_1_ (Fig 4G-H). These results suggest that, while the γ_1_ subunit fails to suppress the current of the embryonic Ca_V_1.1e splice variant by reducing its amplitude (Fig. 3A-C), it still inhibits Ca_V_1.1e currents like in Ca_V_1.1a, by left-shifting the steady state inactivation and causing more complete inactivation, thus reducing the window current.

The γ_1_ subunit was also reported to accelerate the inactivation kinetics of Ca_V_1.1 (Ahern et al. 2001; Freise et al. 2000). Accordingly, the time constant of the slow component of inactivation of Ca_V_1.1a was significantly reduced in the presence of γ_1_ (Fig. S1E-H, Table S2). On the other hand, the time constant of the slow component of inactivation of the embryonic isoform Ca_V_1.1e is already significantly faster than that of the adult Ca_V_1.1a isoform (Ca_V_1.1e: 2.9 s; Ca_V_1.1: 7.8 s), as previously reported (Tuluc et al. 2009). Co-expression of γ_1_ further accelerates the kinetics of inactivation of Ca_V_1.1e, although not to a statistically significant extent (Fig. S1E-H and Table S2).

### The γ_1_ subunit increases membrane expression of both Ca_V_1.1 isoforms

Ca_V_1.1 is the only one out of the ten voltage-gated calcium channels that expresses poorly in non-muscle cells, unless the adaptor protein STAC3 is co-expressed (Polster et al. 2015). Recently it was shown that also the γ_1_ subunit supports robust membrane expression of Ca_V_1.1a in tsA201 cells; although in the absence of STAC3 these channels produce only very small calcium currents (Polster et al. 2016). To examine whether the γ_1_ subunit supports only the membrane targeting of the adult Ca_V_1.1a isoform or also of the embryonic Ca_V_1.1e, we established a dual-labelling approach, originally developed by the lab of Henry Colecraft (Fang and Colecraft 2011; Yang et al. 2010), to identify and quantify membrane inserted Ca_V_1.1 channels. To this end, a 13 amino acid high affinity bungarotoxin (BTX) binding site (BBS) was introduced into the extracellular IIS5-IIS6 domain of GFP-Ca_V_1.1a and GFP-Ca_V_1.1e. Then the channels expressed on the cell surface of HEK cells (expressing β_3_ and α_2_δ-1) were labeled by exposing non-permeabilized living cells to biotinylated bungarotoxin and subsequently to streptavidin-coated quantum dots (QD_655_) (Fig. 5A). Hence, the GFP fluorescence of a cell measures the total Ca_V_1.1 expression, while the QD_655_ fluorescence quantifies the fraction of surface-expressed Ca_V_1.1 channels.

**Figure 5.**
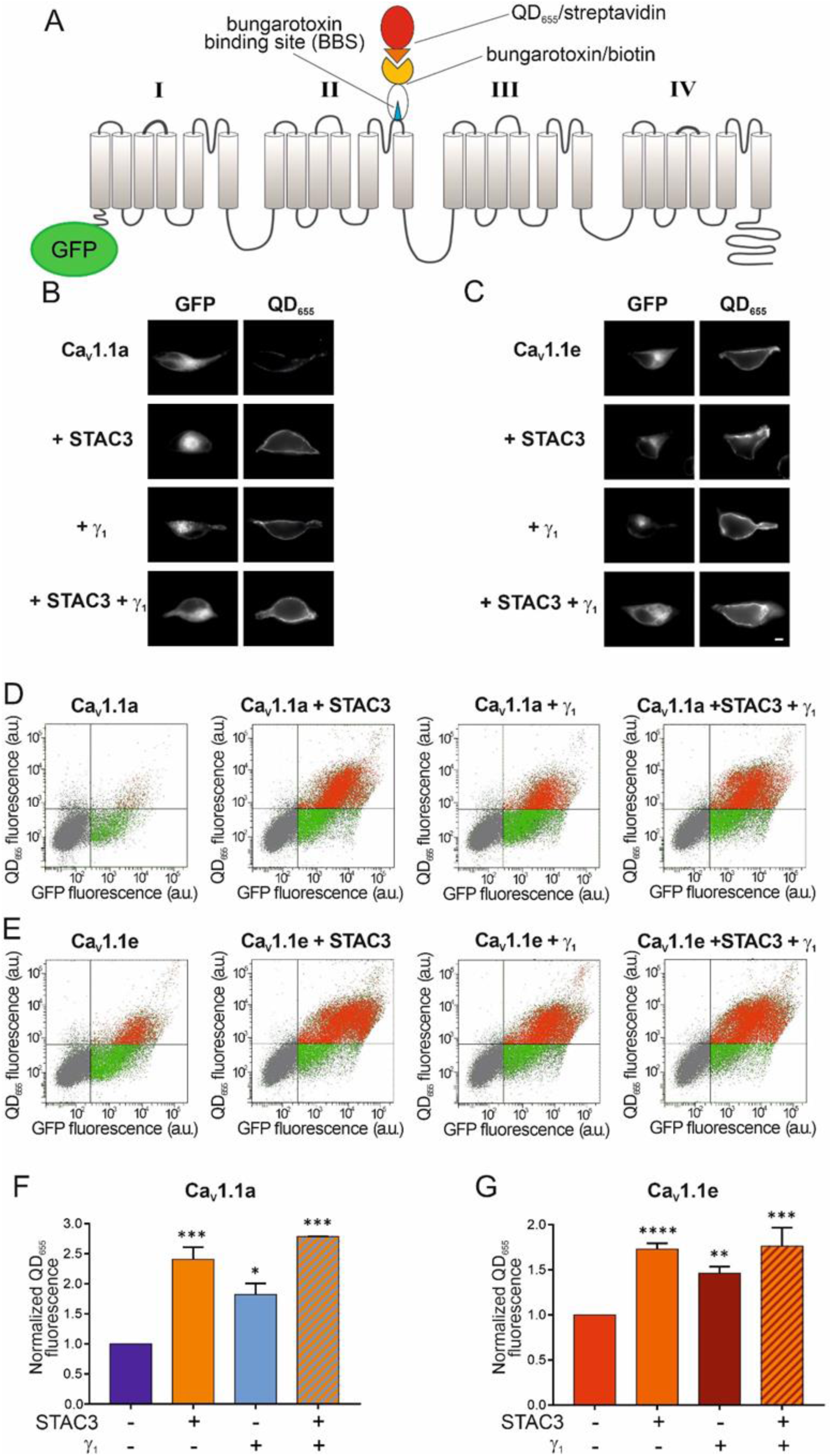
γ_1_ increases the surface density of both Ca_V_1.1a and Ca_V_1.1e isoforms. **(A)** Scheme displaying the strategy to detect Ca_V_1.1 channels expressed on the plasma membrane of HEK cells (stably expressing β_3_ and α_2_δ-1). The introduction of the 13 amino acid BBS in the extracellular domain of GFP-Ca_V_1.1a or GFP-Ca_V_1.1e allowed the selective labelling of channels in the membrane by sequentially incubating the non-permeabilized cells with biotinylated bungarotoxin and streptavidin-conjugated quantum dots (QD_655_). **(B)** From top to bottom, representative images of HEK cells expressing the adult GFP-Ca_V_1.1a isoform alone, with STAC3, with γ_1_, and with both STAC3 and γ_1_. **(C)**. The same for HEK cells expressing the embryonic GFP-Ca_V_1.1e isoform. Scale bar, 2 µm. **(D-E)** Representative raw data from flow cytometry experiments showing the GFP and the QD_655_ signal for cells expressing GFP-Ca_V_1.1a **(D)** or GFP-Ca_V_1.1e **(E)** alone, with STAC3, with γ_1_, and with both STAC3 and γ_1_. The vertical and horizontal lines represent threshold values determined using untransfected cells, untreated cells, and cells exposed only to QD_655_. Single cells are depicted as dots, which have been colored in grey (untransfected), in green (transfected,lacking surface expression) or in red (transfected,with appreciable surface expression). **(F-G)** Normalized mean QD_655_ fluorescence signals across separate flow cytometry experiments (N=4). Data were normalized to the QD_655_ signals of cells expressing only GFP-Ca_V_1.1. In **(F)** the conditions with STAC3 (***, p=0.0003), γ_1_ (*, p=0.0143), and STAC3 + γ_1_ (***, p=0.0002) are significantly different from the control GFP-Ca_V_1.1a using one-way ANOVA and Tukey post hoc mean comparison. In **(G)** the conditions with STAC3 (****, p<0.0001), γ_1_ (**, p=0.0019), and STAC3 + γ_1_ (***, p=0.0002) are significantly different from the control GFP-Ca_V_1.1e using one-way ANOVA and Tukey post hoc mean comparison.

In cells expressing Ca_V_1.1a alone, we detected minimal QD_655_ fluorescence in the plasma membrane. By contrast, co-expression of STAC3 or γ_1_, individually or together, all resulted in robust Ca_V_1.1a membrane targeting (Fig. 5B). In order to quantify membrane inserted Ca_V_1.1 channels, we used flow cytometry analysis, which allows measuring the fluorescence signals of a multitude of individual cells (Fig. 5D). This analysis confirmed the lack of a robust QD_655_ fluorescence signal in cells expressing only GFP-Ca_V_1.1a, but displayed strong QD_655_ fluorescence in cells co-expressing GFP-Ca_V_1.1a together with STAC3, γ_1_, or both. In four independent experiments, cells co-expressing STAC3 on average displayed a 140% increase of surface expressed Ca_V_1.1a, cells co-expressing γ_1_ an 80% increase, while cells expressing both, STAC3 and γ_1_ subunits, displayed a 180% increase compared to cells expressing Ca_V_1.1a alone (Fig. 5F). These results corroborate the importance of STAC3 and γ_1_ for Ca_V_1.1a plasma membrane expression (Niu et al. 2018; Polster et al. 2016; Polster et al. 2015).

We then analyzed the effect of the STAC3 and γ_1_ subunits on membrane expression of the embryonic Ca_V_1.1e isoform. In contrast to the adult isoform, the embryonic Ca_V_1.1e channel showed a substantial membrane staining also when expressed alone (Fig. 5C top, 5E left). Nevertheless, co-expression of STAC3 and γ_1_, individually or together, further increased the amount of QD_655_ fluorescence (Fig. 5C, 5E). In four independent experiments, cells co-expressing STAC3 displayed a 70% increase of surface-expressed Ca_V_1.1e, cells co-expressing γ_1_ a 50% increase, while the ones expressing both, STAC3 and γ_1_ subunits, displayed an 80% increase compared to cells expressing Ca_V_1.1e alone (Fig. 5G).

All together, these results demonstrate that, while the γ_1_ subunit fails to modulate the current amplitude of the embryonic Ca_V_1.1e isoform, it still modulates its steady-state inactivation and surface trafficking. Moreover, the reduction of current density induced by γ_1_ cannot be explained by reduced channel availability at the cell surface.

### Ca_V_1.1–γ_1_ ion-pair partners predicted by structure modeling are not essential for Ca_V_1.1a-specific current reduction by γ_1_

Since the recent cryo-EM structure of Ca_V_1.1 revealed that the γ_1_ subunit interacts with IVS3-S4 (Wu et al. 2016; Wu et al. 2015) and because we found that γ_1_ fails to inhibit the current amplitude of the embryonic Ca_V_1.1e isoform (Fig. 2E), which lacks 19 amino acids in the IVS3-S4 linker, we hypothesized that γ_1_ and the IVS3-S4 linker of Ca_V_1.1a may establish an interaction responsible for the current inhibition in Ca_V_1.1a. In order to identify putative interaction partners between the IVS3-S4 linker and γ_1_, we generated a structural model of the Ca_V_1.1 channel based on the published cryo-EM structure (Wu et al. 2016) (Fig. S2). We used the Rosetta computational modeling software (Bender et al. 2016; Rohl et al. 2004) to model the structure of the IVS3-S4 linker of Ca_V_1.1a. The resulting structure predicts a putative interaction of residues D1223 and D1225 of the IVS3-S4 linker of Ca_V_1.1a with residue R160 in the second extracellular loop of the γ_1_ subunit (Fig. 6A, Fig. S2). To test whether the observed inhibition of the Ca_V_1.1a current amplitude by γ_1_ is dependent on this ionic interaction, we performed site-directed mutagenesis to substitute the involved residues with alanines, which deletes all interactions made by side-chain atoms beyond the β carbon (Wells 1991). However, mutation of residue R160 of the γ_1_ subunit to an alanine did not diminish its ability to inhibit the current amplitude of Ca_V_1.1a (Fig. 6A-D, Table S3). Also simultaneously mutating both D1223 and D1225 of Ca_V_1.1a did not alter the ability of γ_1_ to reduce the current amplitude of Ca_V_1.1a (Fig. 6E-H, Table S3). Together these results indicate that this putative interaction between the IVS3-S4 linker of Ca_V_1.1a and the γ_1_ subunit is dispensable for current amplitude inhibition by γ_1_.

**Figure 6.**
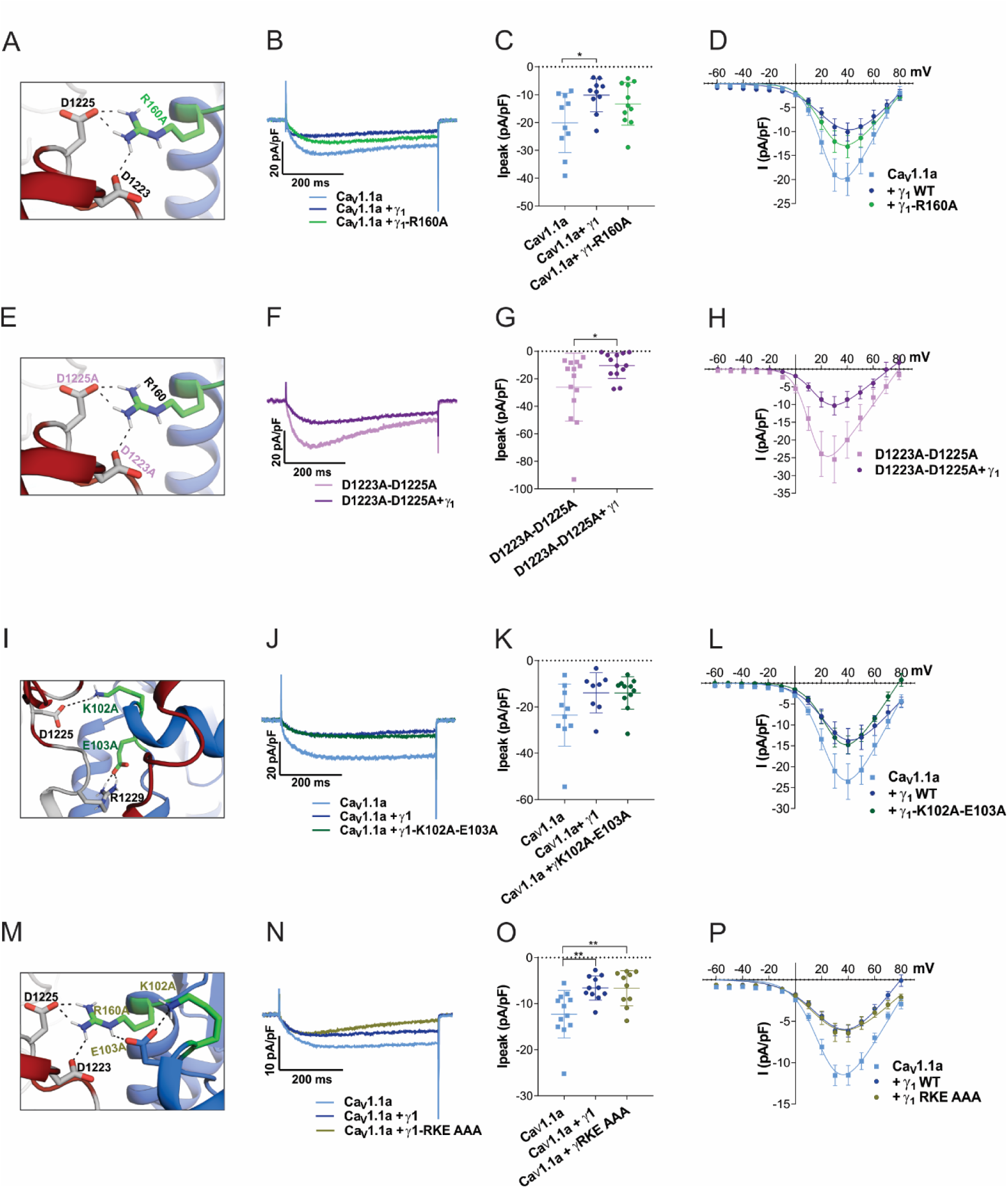
The putative interactions between the IVS3-S4 loop and γ_1_ identified by structure modeling are dispensable for Ca_V_1.1a current reduction. **(A-H)** Structure modelling of Ca_V_1.1a and γ_1_ indicates interactions of R160 (γ_1_) with D1223 and D1225 (Ca_V_1.1a). Neutralizing the putative γ_1_ interaction partner (R160A) **(A)** or the Ca_V_1.1a interaction partners (D1223A and D1225A) **(E)**, did not impair current reduction by γ_1_ **(B-D, F-H). (I-L)** Structure modelling of Ca_V_1.1a and γ_1_ indicates further interactions of K102 and e103 (γ_1_) with D1225 and R1229 (Ca_V_1.1a). Neutralizing both of these putative Ca_V_1.1a interaction partners to alanine (K102A/E103A) **(I)** did not abolish the ability of γ_1_ to reduce Ca_V_1.1a current **(J-K)**. Also concomitant mutation of all three γ_1_ residues involved in these putative interactions did not abolish the current reduction by γ_1_ **(N-P). (B, F, J, N)** Exemplary current traces at V_max_, **(C, G, K, O)** scatter plots of I_peak_ and **(D, H, L, P)** current-voltage relationship. Mean±SEM; P-values calculated with ANOVA and Tukey’s post hoc test. * *P<*0.05 and ** P<0.01.

Previously it has been suggested that the N-terminal half of the γ_1_ subunit, including the first two transmembrane domains, mediates its interaction with the calcium channel and is responsible for suppressing the current amplitude of Ca_V_1.1 (Arikkath et al. 2003). Because the analyzed R160A mutation is located in the C-terminal half of the γ_1_ subunit protein, we modeled the structure of the extensive extracellular loop located in the first half of the γ_1_ subunit and searched it for further possible interaction sites. We identified putative ionic interactions of residues D1225 and R1229 in the IVS3-S4 linker of Ca_V_1.1a with the K102 and E103 positioned in the first extracellular domain of the γ_1_ subunit (Fig. 6I, Table S3, Fig. S2). However, mutation of K102 and E103 to alanines did not alter the ability of γ_1_ to inhibit the calcium channel current amplitude (Fig. 6J-L, Table S3). Finally, to exclude the possibility that the interaction between the IVS3-S4 linker of Ca_V_1.1a with either one of the two extracellular loops of γ_1_ were sufficient to suppress the calcium channel current amplitude, we combined the R160A and the K102A/E103A mutations (Fig. 6M). However, also this triple-mutant γ_1_ was capable of inhibiting the current amplitude of Ca_V_1.1a to similar levels as the wild-type γ_1_ (Fig. 6N-P, Table S3). Together these mutagenesis experiments indicate that the current-inhibiting effect of γ_1_ is not mediated by direct ionic interactions between γ_1_ and the IVS3-S4 loop of Ca_V_1.1a.

## Discussion

Whereas the role of the auxiliary α_2_δ and β subunits in subcellular targeting and gating modulation have been extensively studied for high-voltage activated Ca^2+^ channels in heterologous cells, this has not been the case for the γ_1_ subunit. γ_1_ is a specific subunit of the skeletal muscle Ca_V_1.1 isoform and, until recently, Ca_V_1.1 had resisted efficient functional expression in heterologous expression systems. Only since the discovery of STAC3 as an essential component of the Ca_V_1.1 channel complex permitting the reliable heterologous expression of Ca_V_1.1 such analyses are possible (Horstick et al. 2013; Nelson et al. 2013; Polster et al. 2015). Here we developed and validated two HEK cell lines stably expressing STAC3 (plus α_2_δ-1 and β_3_), which proved to be a convenient and efficient heterologous expression system for Ca_V_1.1. By co-expression of Ca_V_1.1 and γ_1_ in these cells, we found three effects of the γ_1_ subunit: facilitated membrane expression, a reduction of the current density, and a shift of steady-state inactivation to hyperpolarizing potentials. The effects of the γ_1_ subunit on the two splice variants of Ca_V_1.1 expressed in our new STAC3-HEK cell lines revealed a novel, isoform-dependent mechanism of channel modulation by this subunit. Although γ_1_ supports membrane expression of Ca_V_1.1a and Ca_V_1.1e, it only functions as a negative regulator of the adult of Ca_V_1.1a splice variant. This differential regulation is mediated by the inclusion of the alternatively spliced exon 29 in the extracellular loop connecting helices S3 and S4 in repeat IV, but it does not require the direct ionic interactions between this loop and the γ_1_ subunit. Another novel finding is that in both, the adult and embryonic Ca_V_1.1 splice variant, γ_1_ reduces steady-state inward current at more negative voltages by shifting the voltage-dependence of steady-state inactivation but not of activation to more negative voltages and by promoting the time course of current inactivation.

### The γ_1_ subunit supports membrane expression of Ca_V_1.1

The substantially increased surface expression induced by co-expression of γ_1_ observed with extracellular bungarotoxin labeling and flow-cytometry did not translate into increased current densities. This is consistent with the observation that in γ_1_-null mouse muscle, in which STAC3 is still endogenously expressed, the expression levels of Ca_V_1.1 are similar to those of wild type mice (Arikkath et al. 2003). In our experiments this is explained by the observation, that the effects of γ_1_ and STAC3 on membrane expression are not additive and therefore γ_1_ does not significantly increase Ca_V_1.1 beyond the level already achieved by STAC3. Apparently, an independent component must be limiting for membrane targeting. The effect of γ_1_ on membrane targeting in heterologous cells is consistent with a previous immunocytochemistry and charge movement analysis showing that in the absence of STAC3, the γ_1_ subunit supports robust membrane expression of Ca_V_1.1 in tsA201 cells, while promoting only very small currents (Polster et al. 2016). On the contrary, an earlier Western blot analysis of tsA201 cells lysates reported that co-expression of γ_1_ reduces the levels of Ca_V_1.1 protein expression (Sandoval et al. 2007). In sum, our results corroborate the findings that the γ_1_ subunit supports membrane expression of Ca_V_1.1 in heterologous cell systems in a splice variant independent manner, possibly by masking retention motives on the C-terminus (Niu et al. 2018); however, without adding to the calcium influx.

### The γ_1_subunit promotes steady-state inactivation in Ca_V_1.1a and Ca_V_1.1e

Functionally, the two negative actions of γ_1_ on Ca_V_1.1 currents dominate. The observed decrease in current amplitude and left-shift of steady-state inactivation are in general agreement with previous studies in muscle cells (Ahern et al. 2001; Freise et al. 2000) as well as in tsA201 cells expressing Cav1.1a (Polster et al. 2016). Limiting calcium influx through Ca_V_1.1 during muscle activity is tolerable because of the principal role of Ca_V_1.1 as voltage sensor in skeletal muscle EC coupling (Schneider and Chandler 1973; Rios and Brum 1987). At the same time, it is important to limit interference of calcium influx with other calcium signaling events, like those regulating fiber type specification, and to avoid adverse effects of calcium overload on the mitochondrial integrity (Sultana et al. 2016). Previously, we pointed out, how intrinsic mechanisms in the Ca_V_1.1 α_1S_ subunit and the actions of auxiliary subunits cooperate in limiting the calcium currents in skeletal muscle (Tuluc et al. 2009; Flucher et al. 2005). Whereas the α_2_δ-1 subunit slows down the activation, the γ_1_ subunit promotes voltage-dependent inactivation at more negative voltages and makes inactivation more complete. This effect was equally observed in the adult and, as shown here for the first time, also in the embryonic splice variant. Together with the observed increase in membrane targeting, this is the first experimental evidence demonstrating that the γ_1_ subunit functionally interacts with the embryonic splice variant Ca_V_1.1e. Therefore this modulatory effect independent of the length of the extracellular loop connecting helices IVS3 and IVS4.

### The γ_1_ subunit reduces the current amplitude specifically in Ca_V_1.1a

The most interesting finding of this study is the differential down-regulation of calcium currents in Ca_V_1.1a vs. Ca_V_1.1e. The small current size is one of the hallmarks of skeletal muscle calcium currents. Our results demonstrate that the γ_1_ subunit is a major determinant of this reduced current density. Whereas in skeletal muscle the adult and embryonic Ca_V_1.1 splice variants differ substantially in voltage-dependence of current activation and in current size, the currents recorded in the HEK cells (stably expressing α_2_δ-1, β_3_, and STAC3) reproduced the difference in V_½_ of activation, but not in current density. Apparently, this difference was due to the lack of one or more muscle-specific factors in the heterologous expression system. As co-expression of γ_1_ restored the reduced current density in Ca_V_1.1a compared to Ca_V_1.1e, the γ_1_ subunit is such a factor. Quantitatively, the difference in current density between the two splice variants was still smaller than that observed when the same constructs were expressed in dysgenic myotubes (Tuluc et al. 2016; Tuluc et al. 2009). Therefore, it is likely that other modulatory mechanisms present in the native environment of the channel in the skeletal muscle triads contribute to expression of this splice variant-specific difference. The γ_1_ subunit is the second identified protein that modulates differently the current properties of the two Ca_V_1.1 splice variants, after the RyR1 (Benedetti et al. 2015), and demonstrates the importance of the native cellular environment for the accurate expression of physiological current properties. Notably, γ_1_ does not reduce the current density of Ca_V_1.1a by decreasing its plasma membrane expression. As previously shown, Ca_V_1.1e has a higher open probability than Ca_V_1.1a in skeletal myotubes (Tuluc et al. 2009). Therefore, the most likely explanation is that γ_1_ decreases the channel’s maximal open probability in a splice variant-specific manner.

The sole difference in the primary structure between the embryonic and adult splice variants is the inclusion of 19 amino acids coded in exon 29 into the IVS3-S4 loop of Ca_V_1.1a. Apparently, this difference determines the action of the γ_1_ subunit on current size. There are two possible mechanisms how inclusion of exon 29 can enable this functional interaction with γ_1_. Direct interactions between the IVS3-S4 loop and γ_1_, or the stabilization of a conformation of the channel complex by inclusion of exon 29 that renders Ca_V_1.1a susceptible to this particular γ_1_ modulation. As the first possibility is amenable to experimental testing, we examined this possibility by identifying and mutating putative interaction sites on both channel subunits. However, none of these ion pairs seemed to be essential for the current-reducing action of γ_1_. Therefore, it is very unlikely that this effect is mediated by the direct interaction of the γ_1_ subunit with the IVS3-S4 loop, although our experiments do not rule out this possibility. Alternatively, we conclude that insertion of exon 29 into this loop alters the conformation of the channel complex in a way that enables it to respond to the inclusion of γ_1_ with a reduced current density (Fig. 7A). Notably, the left-shifted activation in Ca_V_1.1e compared to Ca_V_1.1a is observed with or without γ_1_, and the left-shifted inactivation is observed with or without exon 29, while the decreased current amplitude requires their co-operation. Evidently, the interdependence of the analyzed gating properties on the IVS3-S4 loop and the γ_1_ subunit is highly specific. Each of the partners independently exerts its specific action on the voltage dependences of activation and inactivation, respectively (Fig. 7B).

**Figure 7.**
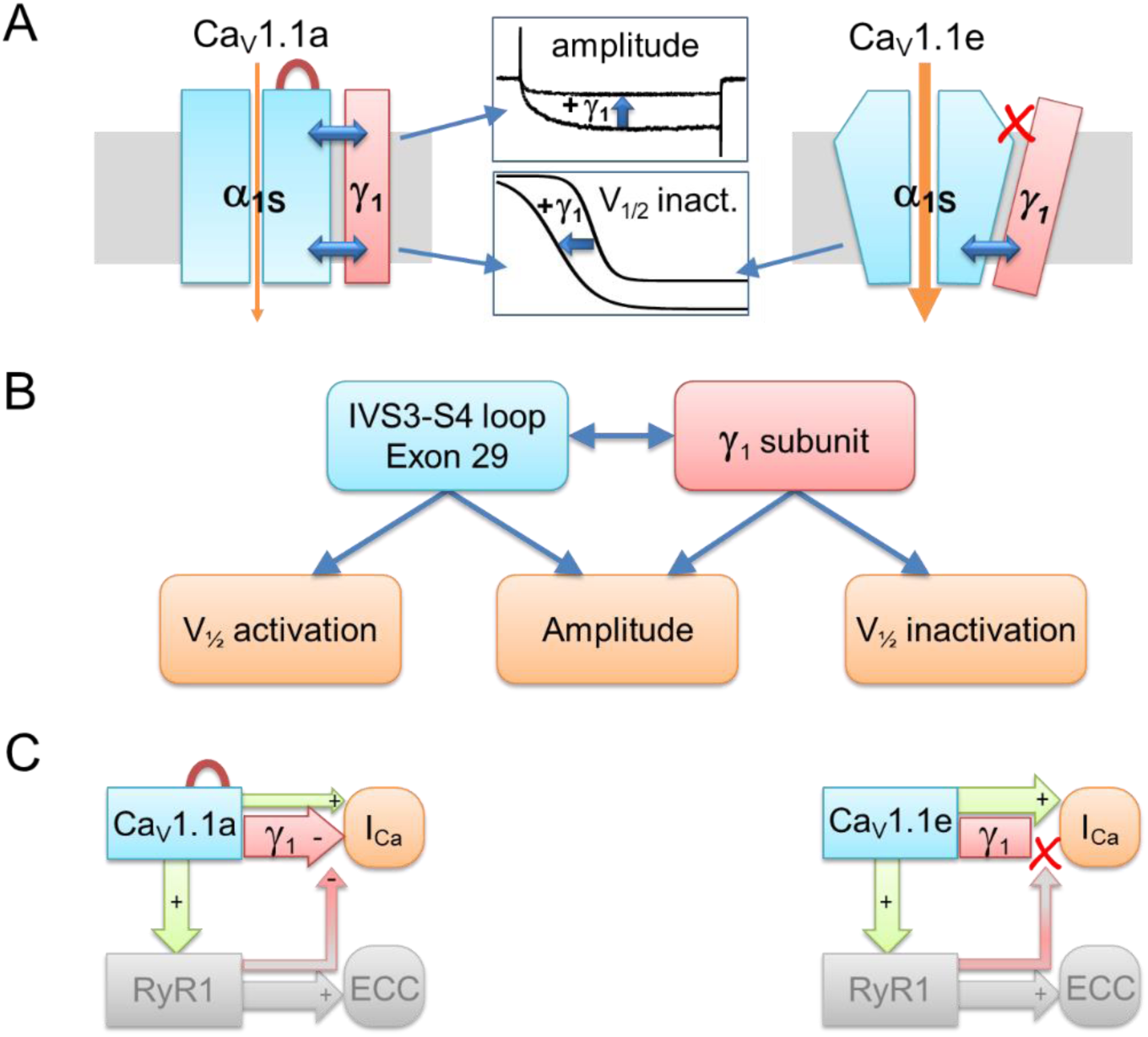
Model of differential γ_1_ modulation on Ca_V_1.1a and Ca_V_1.1e currents and its consequences for retrograde coupling. **(A)** In both Ca_V_1.1 splice variants the γ_1_ subunit limits calcium currents by shifting the voltage-dependence of inactivation to more hyperpolarizing potentials and rendering inactivation more complete. Inclusion of exon 29 in the extracellular IVS3-S4 loop stabilizes a conformation of the Cav1.1a channel complex, which enables the γ_1_ subunit to reduce the current amplitude. **(B)** The IVS3-S4 loop including exon 29 and the γ_1_ subunit require each other for reducing the current amplitude. In contrast, this cooperation is not required to shift the voltage dependence of activation and inactivation, which occurs in a splice-variant dependent manner. **(C)** In skeletal muscle cells, the negative regulation of calcium currents by the γ_1_ subunit is a prerequisite of retrograde current amplification by the RyR1 in Ca_V_1.1a (red arrow from RyR1 to γ_1_) (Grabner et al. 1999; Nakai et al. 1996). Without exon 29 in embryonic Ca_V_1.1e, no γ_1_-dependent reduction of current amplitude and no RyR1-dependent relief of this inhibition occurs (Benedetti et al. 2015). The red loop in Ca_V_1.1a indicates inclusion of exon 29.

### The role of the γ_1_ subunit in retrograde coupling of Ca_V_1.1 and RyR1

In skeletal muscle Ca_V_1.1a calcium currents are augmented by an interaction of its cytoplasmic II-III loop with the RyR1 (Grabner et al. 1999). Previously we demonstrated that this function, termed retrograde coupling, is specific for the adult Ca_V_1.1a splice variant (Benedetti et al. 2015). The currents of Ca_V_1.1e are not reduced when the connection with the RyR1 is severed. The dependence of the current augmentation by retrograde coupling on inclusion of exon 29 into the IVS3-S4 loop of Ca_V_1.1 mirrors the importance of exon 29 for the current reduction by γ_1_. Based on the results of the earlier study we had proposed a mechanistic model according to which, retrograde coupling partially relieves the inhibition of Ca_V_1.1 currents by an unknown, exon 29-dependent factor. Our current study identifies the γ_1_ subunit as this inhibitory factor. In the simultaneous presence of exon 29 and the γ_1_ subunit, the currents of Ca_V_1.1a are reduced, and this effect is partially counteracted by the interaction with RyR1. If either exon 29 or the γ_1_ subunit are missing, this inhibition is absent and there is nothing to be relieved by retrograde coupling (Fig. 7C).

## Conclusions

This analysis of the actions of the γ_1_ subunit on the two splice variants of Ca_V_1.1 in heterologous cells revealed multiple functions of γ_1_ in membrane targeting and functional modulation of the skeletal muscle calcium channel. Interestingly, some of the γ_1_ effects are general for both splice variants, while another is specific for the adult Ca_V_1.1a. Inclusion of exon 29 in Ca_V_1.1a appears to allosterically render the channel susceptible to the reduction of its currents by γ_1_, as well as to the simultaneous relieve of this block by RyR1. Newly available mammalian cell systems proved highly valuable for this type of co-expression study of Ca_V_1.1, but at the same time highlight the multitude of factors involved in shaping the physiological current properties in its native environment of skeletal muscle.

## Acknowledgements

We thank Katharina Heinz, Sandra Demetz, Irene Mahlrecht, Nicole Kranebitten, Enikō Török and Martin Heitz for excellent technical help. This work was supported by grants from the Tiroler Wissenschaftsfond 2018 (UNI-0404-2238 to MC), and from the Austrian Science Fund (FWF) (T855 and P33776 to MC, P30402 to BEF, P27809 to JS) and the Erika-Cremer habilitation fellowship of the University of Innsbruck to NJO. YEG, MFQ, and WET are students of the Ca_V_X PhD program co-funded by FWF (DOC30) and the Medical University Innsbruck. HJD is an employee of Boehringer Ingelheim Pharma GmbH & Co KG. The authors declare no other competing financial interests.

Author contributions: MC and BEF conceived and designed the study. MC made the constructs, RT-PCR, WB analysis, flow cytometry stainings and acquired images. NJO generated the cell lines and acquired electrophysiological data for the HEK-STAC3 cell line under the supervision of JS. YEG and WET acquired all the remaining electrophysiological data under the supervision of MC, PT and BEF. WP and DW acquired and analyzed the flow cytometry data. MFQ and SM performed the modeling under the supervision of KL. HJD provided the cell line expressing β_3_ and α_2_δ-1. MC, BEF and YEG wrote the manuscript. All authors contributed to the final draft of the manuscript.

## Supplementary figures

**Figure S1.**
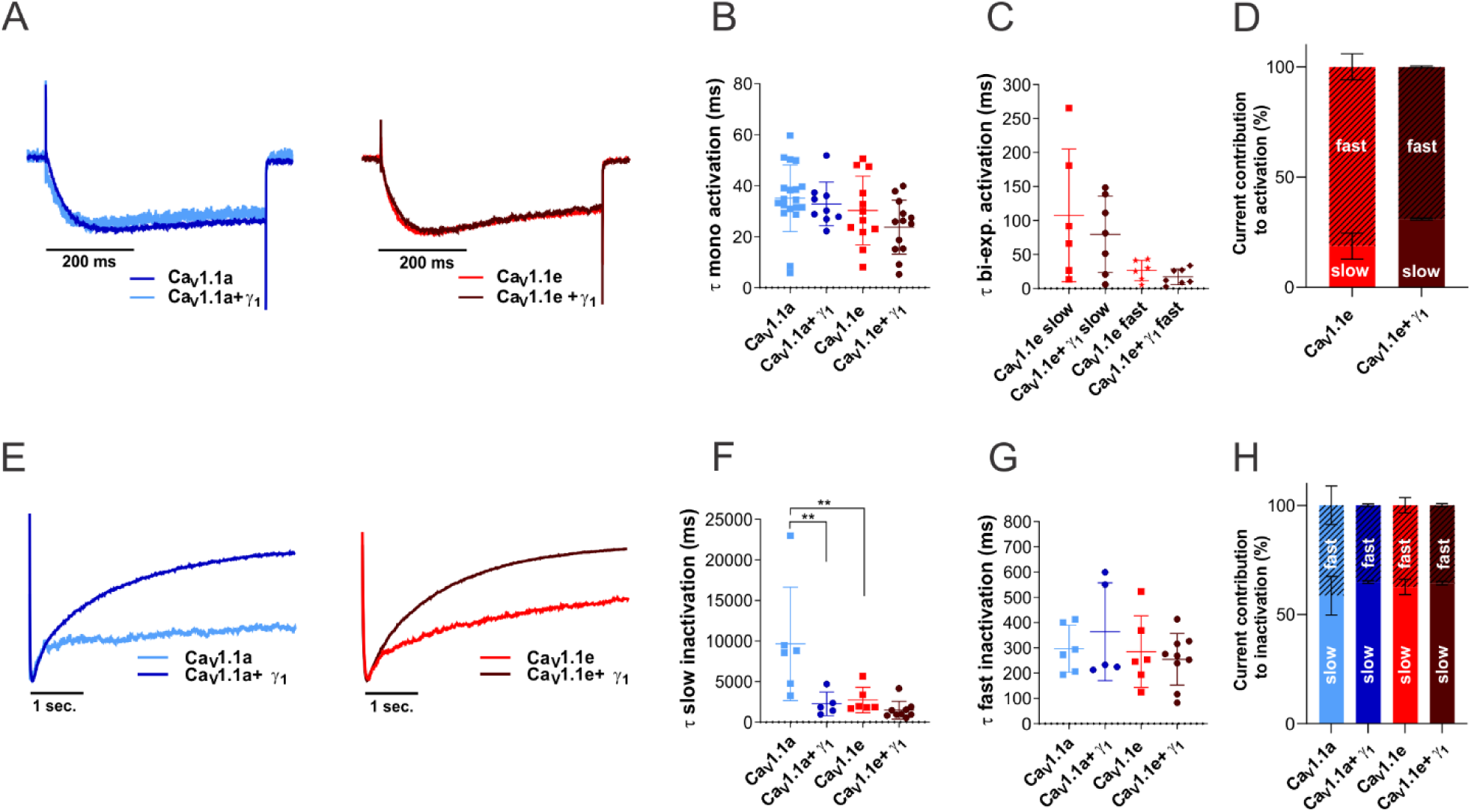
γ_1_ does not affect activation kinetics but accelerates inactivation kinetics in Ca_V_1.1a. **(A-D)** Time constants of activation of Ca_V_1.1a (blue, n=19), Ca_V_1.1a + γ_1_ (dark blue, n=9), Ca_V_1.1e (red, n=12) and Ca_V_1.1e + γ_1_ (dark brown, n=13) of a mono-exponential and bi-exponential fit (Ca_V_1.1e) on the rising phase of the inward calcium current during a 500 ms depolarization to V_max_. **(A)** Example traces of 500 ms depolarization to V_max_ in Ca_V_1.1a (left) and Ca_V_1.1e (right), normalized to the peak current. No differences were found between the time constant of activation of Ca_V_1.1a or Ca_V_1.1e with or without γ_1_ co-expression when fitted mono-exponentially **(B)** or between the fast or slow time constant of Ca_V_1.1e (n=6) and Ca_V_1.1e + γ_1_ (n=7) of the recordings that could be fitted bi-exponentially **(C)**, Ca_V_1.1a and Ca_V_1.1a + γ_1_ could only be fitted mono-exponentially. The current contribution of the fast component was bigger than that of the slow component in both Ca_V_1.1e (slow:fast ≈ 20:80) and Ca_V_1.1e + γ_1_ (slow:fast ≈ 30:70), but the ratios were similar (p=0.41) **(D). (E-H)** Slow and fast time constant of inactivation of Ca_V_1.1a (blue, n=6), Ca_V_1.1a + γ_1_ (dark blue, n=5), Ca_V_1.1e (red, n=6) and Ca_V_1.1e + γ_1_ (dark brown, n=9) of a bi-exponential fit on the decay phase of the inward calcium current during a 5 sec. depolarization to V_max_. **(E)** Example traces of 5 sec. depolarization to V_max_ in Ca_V_1.1a (left) and Ca_V_1.1e (right), normalized to the peak current. A significant acceleration (p=0.007) of the slow time constant of inactivation was found when Ca_V_1.1a was co-expressed with γ_1_, co-expression of Ca_V_1.1e with γ_1_ shows a 2-fold, but not significant, acceleration (p=0.88) **(F)**. No differences were found between Ca_V_1.1a and Ca_V_1.1e with or without γ_1_ co-expression in the fast time constant of inactivation **(F)**. The ratios of current contribution to inactivation of the slow versus fast component were similar between all four groups (p=0.61 for Ca_V_1.1a vs. Ca_V_1.1a + γ_1_, p=0.79 for Ca_V_1.1e vs. Ca_V_1.1e + γ_1_), with a somewhat higher contribution of the slow than the fast component (slow:fast ≈ 60:40). Mean±SEM; Significance was calculated with ANOVA and Sidak’s post hoc test. ** *P*<0.001. P-values for current contributions were calculated with Student’s t-test.

**Figure S2.**
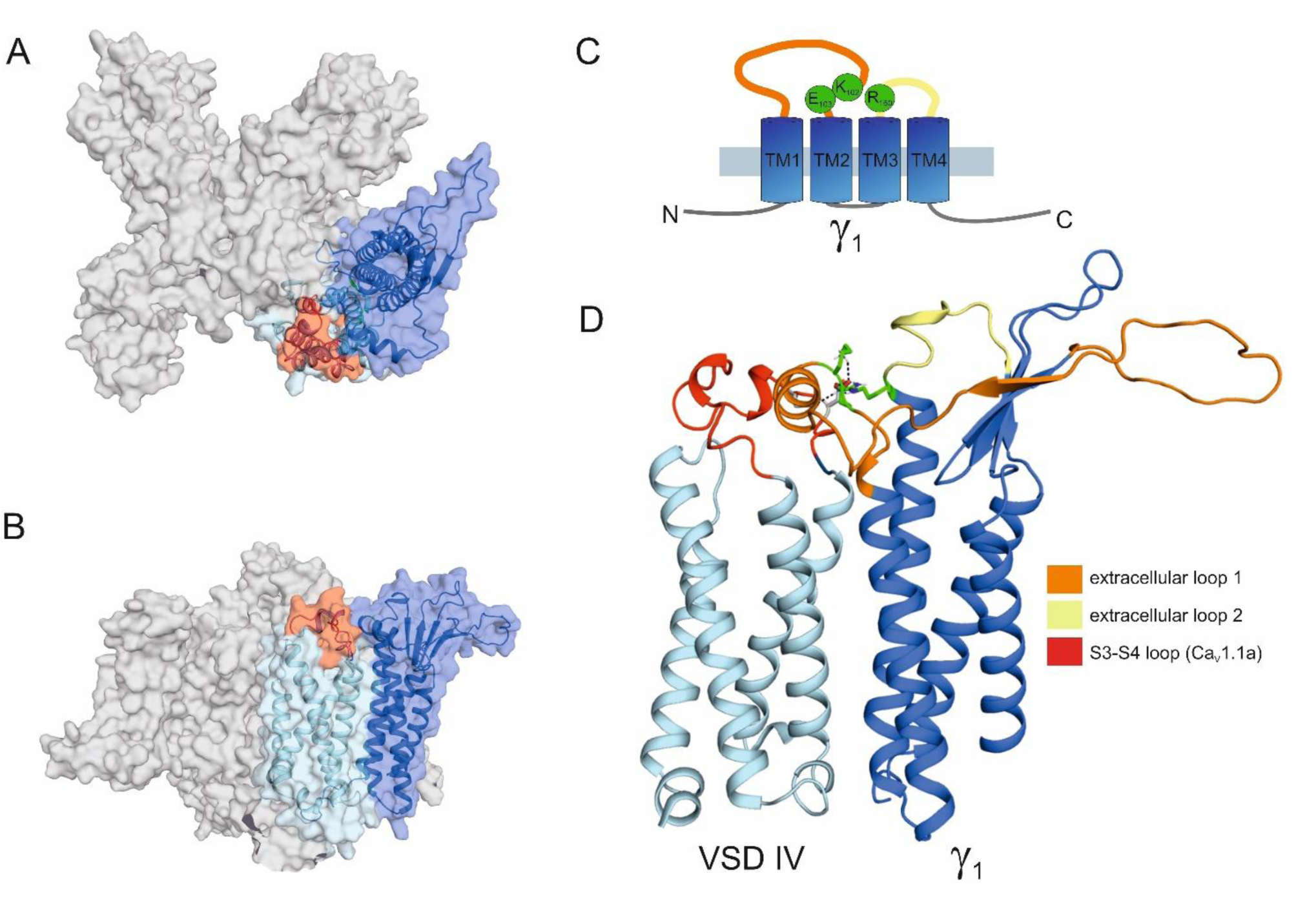
Structure modeling of Ca_V_1.1a in complex with the γ_1_ subunit. (**A**) Top view of the structure model of the human Ca_V_1.1 α_1_ subunit in complex with the γ_1_ subunit refined with molecular dynamics (MD) simulation in a membrane environment based on the 3.6 Å structure of rabbit Ca_V_1.1 (Wu et al. 2016). VSD IV (light blue) is interacting with the γ_1_ subunit (marine). The alternatively spliced exon 29 (red) is inserted in the IVS3-S4 linker of Ca_V_1.1a. (**B**) Side view of the structure model of the Ca_V_1.1 with the γ_1_ subunit. (**C**) Cartoon showing the domain organization of γ_1_, with the mutated residues R160, K102 and E103 in green. (**D**) Close-up of the interaction side of the VSD IV S3-S4 loop with the γ_1_ subunit, highlighting the extracellular loops of the γ_1_ subunit. The extracellular loop 1 of γ_1_ is in the same orientation as presented in Figure 6 (Panels A, E, M) and residue R160 is highlighted in green.

**Table S1.** Current-voltage parameters (activation) for whole-cell electrophysiology experiments and fit equation of Ca_V_1.1a and Ca_V_1.1e in HEK-STAC3 and HEK-TetOn-STAC3

|  | HEK-STAC3 |  |  | HEK-TetOn-STAC3 |  |  |
| --- | --- | --- | --- | --- | --- | --- |
| | $\text{Ca}_v1.1a$ | $\text{Ca}_v1.1e$ | p-value (t-test) | $\text{Ca}_v1.1a$ | $\text{Ca}_v1.1e$ | p-value (t-test) |
| $I_{\text{peak}}$ (pA/pF) | -19.8±2.7 | -19.5±2.4 | 0.94 | -18.2±3.1 | -21.3±4.5 | 0.56 |
| $V_{1/2}$ act. (mV) | 25.1±0.7 | 4.5±1.0 | **** | 24.1±1.4 | 3.0±1.5 | **** |
| k act. (mV) | 8.4±0.3 | 7.2±0.2 | *** | 7.0±0.5 | 5.3±0.3 | 0.011* |
| $V_{\text{rev}}$ (mV) | 84.2±0.9 | 73.3±1.2 | **** | 81.2±1.7 | 72.8±1.6 | 0.0013** |
| time to peak (ms) | 173.1±10.8 | 162.6±12.0 | 0.52 | 198.3±20.4 | 184.1±14.3 | 0.06 |
| n | 15 | 15 | -- | 19 | 15 | -- |

Data are expressed as mean values ± SEM.

**Table S2.**
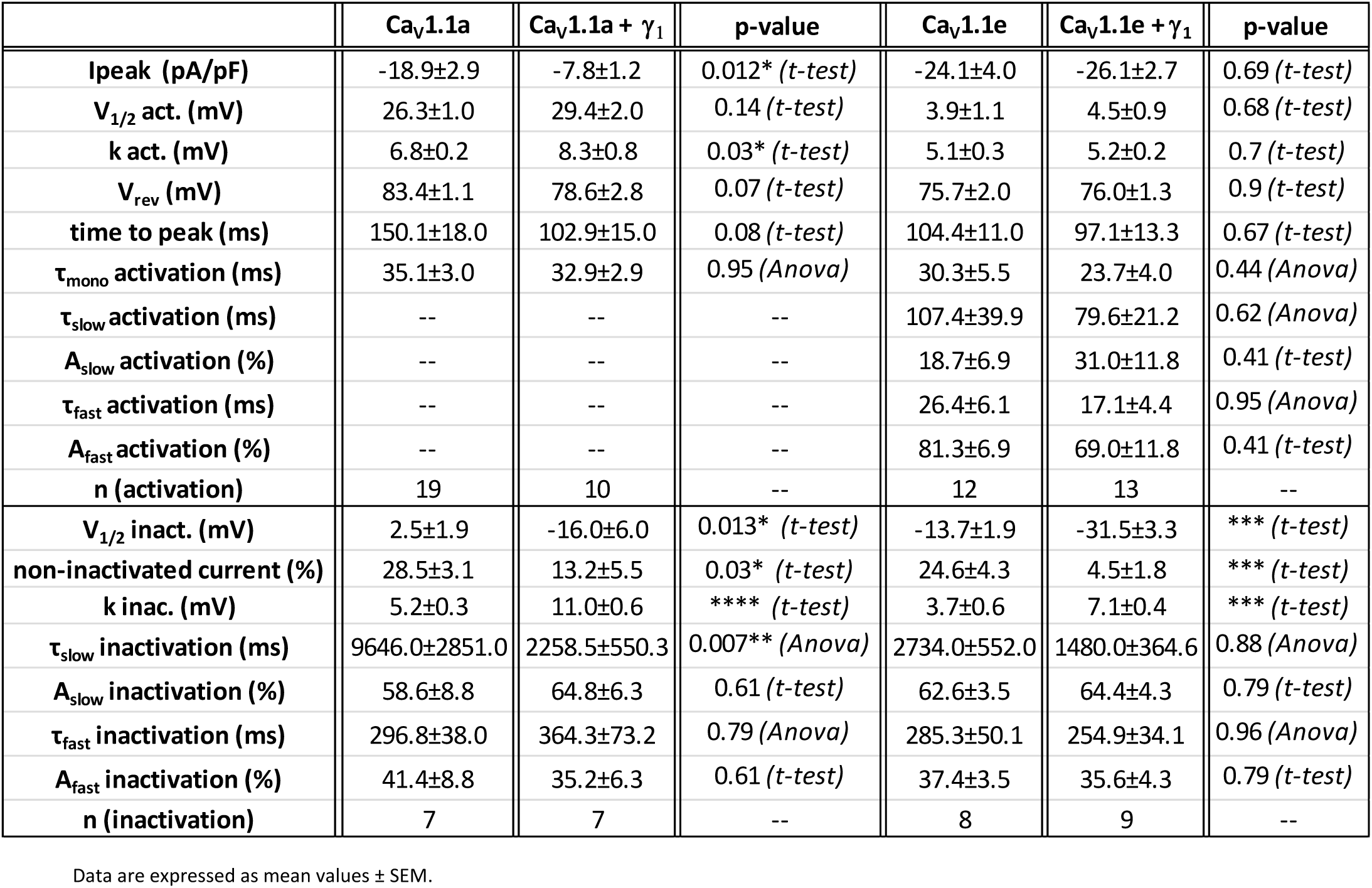
Current-voltage parameters (activation and inactivation) for whole-cell electrophysiology experiments and fit equation of Ca_V_1.1a and Ca_V_1.1e in the presence and absence of γ_1_

| | $\text{Ca}_v1.1a$ | $\text{Ca}_v1.1a + \gamma_1$ | p-value | $\text{Ca}_v1.1e$ | $\text{Ca}_v1.1e + \gamma_1$ | p-value |
| --- | --- | --- | --- | --- | --- | --- |
| $I_{\text{peak}}$ (pA/pF) | -18.9±2.9 | -7.8±1.2 | 0.012* (t-test) | -24.1±4.0 | -26.1±2.7 | 0.69 (t-test) |
| $V_{1/2}$ act. (mV) | 26.3±1.0 | 29.4±2.0 | 0.14 (t-test) | 3.9±1.1 | 4.5±0.9 | 0.68 (t-test) |
| k act. (mV) | 6.8±0.2 | 8.3±0.8 | 0.03* (t-test) | 5.1±0.3 | 5.2±0.2 | 0.7 (t-test) |
| $V_{\text{rev}}$ (mV) | 83.4±1.1 | 78.6±2.8 | 0.07 (t-test) | 75.7±2.0 | 76.0±1.3 | 0.9 (t-test) |
| time to peak (ms) | 150.1±18.0 | 102.9±15.0 | 0.08 (t-test) | 104.4±11.0 | 97.1±13.3 | 0.67 (t-test) |
| $\tau_{\text{mono}}$ activation (ms) | 35.1±3.0 | 32.9±2.9 | 0.95 (Anova) | 30.3±5.5 | 23.7±4.0 | 0.44 (Anova) |
| $\tau_{\text{slow}}$ activation (ms) | -- | -- | -- | 107.4±39.9 | 79.6±21.2 | 0.62 (Anova) |
| $A_{\text{slow}}$ activation (%) | -- | -- | -- | 18.7±6.9 | 31.0±11.8 | 0.41 (t-test) |
| $\tau_{\text{fast}}$ activation (ms) | -- | -- | -- | 26.4±6.1 | 17.1±4.4 | 0.95 (Anova) |
| $A_{\text{fast}}$ activation (%) | -- | -- | -- | 81.3±6.9 | 69.0±11.8 | 0.41 (t-test) |
| n (activation) | 19 | 10 | -- | 12 | 13 | -- |
| $V_{1/2}$ inact. (mV) | 2.5±1.9 | -16.0±6.0 | 0.013* (t-test) | -13.7±1.9 | -31.5±3.3 | *** (t-test) |
| non-inactivated current (%) | 28.5±3.1 | 13.2±5.5 | 0.03* (t-test) | 24.6±4.3 | 4.5±1.8 | *** (t-test) |
| k inac. (mV) | 5.2±0.3 | 11.0±0.6 | **** (t-test) | 3.7±0.6 | 7.1±0.4 | *** (t-test) |
| $\tau_{\text{slow}}$ inactivation (ms) | 9646.0±2851.0 | 2258.5±550.3 | 0.007** (Anova) | 2734.0±552.0 | 1480.0±364.6 | 0.88 (Anova) |
| $A_{\text{slow}}$ inactivation (%) | 58.6±8.8 | 64.8±6.3 | 0.61 (t-test) | 62.6±3.5 | 64.4±4.3 | 0.79 (t-test) |
| $\tau_{\text{fast}}$ inactivation (ms) | 296.8±38.0 | 364.3±73.2 | 0.79 (Anova) | 285.3±50.1 | 254.9±34.1 | 0.96 (Anova) |
| $A_{\text{fast}}$ inactivation (%) | 41.4±8.8 | 35.2±6.3 | 0.61 (t-test) | 37.4±3.5 | 35.6±4.3 | 0.79 (t-test) |
| n (inactivation) | 7 | 7 | -- | 8 | 9 | -- |

Data are expressed as mean values ± SEM.

**Table S3.** Current-voltage parameters (activation) for whole-cell electrophysiology experiments and fit equation of Ca_V_1.1a-D1223A-D1225A, γ_1_-R160A, γ_1_-K102A-E103A and γ_1_-R160A-K102A-E103A (RKEAAA) mutants

| | Ca<sub>v</sub>1.1a | Ca<sub>v</sub>1.1a+ $\gamma_1$ | p-value (ANOVA) | Ca<sub>v</sub>1.1a+ $\gamma_1$-R160 | p-value (ANOVA) |
| --- | --- | --- | --- | --- | --- |
| I<sub>peak</sub> (pA/pF) | -20.1±3.4 | -10.1±1.9 | 0.04* | -13.3±2.3 | 0.20 |
| V<sub>1/2</sub> act. (mV) | 23.1±0.6 | 30.7±2.2 | 0.004** | 24.1±1.3 | 0.88 |
| k act. (mV) | 7.0±0.3 | 13.1±1.8 | 0.001** | 8.5±0.4 | 0.58 |
| V<sub>rev</sub> (mV) | 82.7±2.2 | 86.1±3.9 | 0.65 | 87.4±1.3 | 0.42 |
| time to peak (ms) | 131.8±18.8 | 102.6±8.4 | 0.26 | 95.3±8.9 | 0.12 |
| n | 10 | 10 | -- | 11 | -- |
| | Ca<sub>v</sub>1.1a D1223A-D1225A | Ca<sub>v</sub>1.1a D1223A-D1225A+ $\gamma_1$ | p-value (t-test) | | |
| I<sub>peak</sub> (pA/pF) | -26.1±6.6 | -10.4±2.6 | 0.04* | -- | -- |
| V<sub>1/2</sub> act. (mV) | 12.8±0.8 | 15.8±0.9 | 0.02* | -- | -- |
| k act. (mV) | 6.3±0.4 | 7.1±0.5 | 0.22 | -- | -- |
| V<sub>rev</sub> (mV) | 73.5±2.5 | 69.7±4.3 | 0.44 | -- | -- |
| time to peak (ms) | 87.2±11.0 | 66.6±10.7 | 0.20 | -- | -- |
| n | 14 | 13 | -- | -- | -- |
| | Ca<sub>v</sub>1.1a | Ca<sub>v</sub>1.1a+ $\gamma_1$ | p-value (ANOVA) | Ca<sub>v</sub>1.1a+ $\gamma_1$-K103A-E104A | p-value (ANOVA) |
| I<sub>peak</sub> (pA/pF) | -23.6±4.2 | -13.9±3.1 | 0.13 | -14.0±2.1 | 0.09 |
| V<sub>1/2</sub> act. (mV) | 25.1±1.0 | 26.9±0.2 | 0.40 | 28.2±0.9 | 0.06 |
| k act. (mV) | 8.7±0.5 | 9.8±0.2 | 0.20 | 8.7±0.4 | 0.99 |
| V<sub>rev</sub> (mV) | 86.7±1.8 | 91.8±0.2 | 0.24 | 81.2±2.7 | 0.17 |
| time to peak (ms) | 121.5±18.5 | 99.8±0.9 | 0.69 | 108.5±13.3 | 0.85 |
| n | 10 | 8 | -- | 11 | -- |
| | Ca<sub>v</sub>1.1a | Ca<sub>v</sub>1.1a+ $\gamma_1$ | p-value (ANOVA) | Ca<sub>v</sub>1.1a+ $\gamma_1$-RKE AAA | p-value (ANOVA) |
| I<sub>peak</sub> (pA/pF) | -12.3±1.5 | -6.6±0.8 | 0.006** | -6.6±1.2 | 0.008** |
| V<sub>1/2</sub> act. (mV) | 20.6±0.9 | 28.2±3.3 | 0.13 | 25.7±4.2 | 0.38 |
| k act. (mV) | 8.9±0.6 | 13.9±2.2 | 0.07 | 13.4±2.0 | 0.12 |
| V<sub>rev</sub> (mV) | 89.8±2.0 | 83.2±4.2 | 0.24 | 92.5±2.9 | 0.76 |
| time to peak (ms) | -- | -- | -- | -- | -- |
| n | 12 | 11 | -- | 10 | -- |

Data are expressed as mean values ± SEM.

